# Stem cell basis of a host driven transmission of antigen packed aerosols: a novel mechanism of natural vaccination for tuberculosis

**DOI:** 10.1101/2020.11.14.382572

**Authors:** Bikul Das, Lekhika Pathak, Sukanya Gayan, Bidisha Pal, Parthajyoti Saikia, Tutumoni Baishya, Nihar Ranjan Das, Rupam Das, Mallika Maral, Ranjit Mahanta, Seema Bhuyan, Pratibha Gautam, Joyeeta Talukdar, Sorra Sandhya, Deepjyoti Kalita, Vijay Swami, Krishna Ram Das, Dayal Krishna Bora, Jagat Ghora, Ista Pulu

## Abstract

Natural vaccination against pathogens are known to be achieved by herd-immunity i.e. infected human host provide immunity to the community by spreading the pathogen. Whether, infected human hosts transmit vesicle packed aerosols of pathogen’s antigen for natural vaccination of the community has not yet been considered. We have explored a traditional healing method of aerosol-inoculation against small pox and tuberculosis in the Sualkuchi-Hajo cultural complex of Kamarupa, an ancient Indian region known for tantra-based healing and spirituality. In the aerosol-inoculation method against TB, selected persons with TB (later identified as smear negative TB subject) are encouraged to spread good nigudah in the community by Kirtan chanting; the good Nigudah are thought to be present within bad-nigudah or invisible krimis (tiny flesh eating living being mentioned in ancient India’s medicinal text Caraka Samhita and Atharva Veda). A 15-years of contact TB investigation study, as well as laboratory study of aerosol obtained from smear negative PTB (SN-PTB) subjects led to the identification of good Nigudah as extracellular vesicles (EVs) filled with *Mtb*-antigen ESAT-6. We then developed a mouse model of aerosol-inoculation using SN-PTB subject derived aerosol EVs, and identified *Mtb* infected mesenchymal stem cells (MSCs) of the lung as the putative source of the ESAT-6+ EVs. These *Mtb* infected MSCs reprogram to altruistic stem cell (ASC) phenotype, which then secrete ESAT-6+ EVs to the aerosols; healthy mice receiving the aerosol develop *Mtb* specific herd immunity. These results expedite our ongoing work on the innate defense mechanism of ASCs against pathogen, and provide a novel mechanism of natural vaccination, where the host extracts appropriate antigens from a pathogen, and then spread it in the community via aerosols.

## INTRODUCTION

Pulmonary tuberculosis (PTB) is a global health burden caused by a rod shaped bacteria, the *Mycobacterium tuberculosis* (*Mtb*). The bacteria spread through aerosol in the community and now infect more than 2 billion people, killing 2 million people yearly. The pathogen enters into the respiratory tract and initiates a host/pathogen interaction with macrophages, an immune cell capable of antigen-delivery and export to other immune cells. However, bacteria may hide away from immune cell for years in a dormant state; we found that *Mtb* hijacks bone marrow (BM) stem cells as their protective niche (1–3), confirmed by others (4–7). Under favorable condition, dormant *Mtb* reactivates in the lung causing extensive tissue damage and cavity formation, leading to clinical diseases of PTB (8–10). Only in 60% of PTB subjects, bacteria can be detected in the sputum and these smear positive cases can be isolated and treated to prevent community transmission. Whereas the rest of PTB cases may continue to spread the pathogen as the bacteria can’t be detected in the sputum even after performing culture or testing for bacterial DNA (11). These smear negative PTB subjects are public health challenge especially in rural health set up, where many of them may escape diagnosis and treatment for years and thus, spread the pathogen in the community. The role of these smear negative PTB subjects in either spreading or preventing the disease in the community is not yet clearly known. While managing PTB subjects in rural India through KaviKrishna Telemedicine Care (12, 13), we have uncovered an Indigenous Knowledge System (IKS) based aerosol-inoculation practice that may provide novel insight about the role of these smear negative PTB (SN-PTB) subjects in spreading herd-immunity, a key element of social medicine approach against the pathogen.

IKS is the knowledge system inherited by indigenous culture, and it operates at local level of interaction through tradition/culture, and exhibit emergent property i.e. capable of producing new knowledge through interaction, and adaption to local environments (14, 15). IKS based tool has been applied to develop effective public health approaches (13, 14, 16). One prominent and ancient example IKS based public health approach could be the JivaUpakarTantra (JUT), also known as Vedic Altruism, a nearly extinct knowledge system of ancient Kamrupa (North East India, part of Bangladhesh and West Bengal) cultural complex that survived in rural pockets of Sualkuchi-Hajo area of Assam, India (17) (supplementary note 1). Our ongoing research indicate that JUT might have been involved in the traditional treatment and bioethical management for small pox in the Sualkuchi-Hajo area during 16-19^th^ century, when traditional inoculation/spiritual healing practice were widely used in Bengal and Assam (18–21). In fact, smallpox inoculation practice may have spread to China and medieval Europe from India through Buddhist monasteries (22, 23). Interestingly, our ongoing IKS related study suggest that the JivaUpakarTantra (JUT) based approach against smallpox was unique, as it was aerosol-based and the underlying principle was Nigudah Jiva, which are considered as the lowest form of life in Jainism (24) as well as ancient Kamarupa tantra (17) (Chapter 1.3 and 1.4, pages 6-19; Reference 17). Accordingly, selected person having smallpox were encouraged by tantrik (traditional healers of ancient Kamarupa) to chant mantras in front of others to transmit their good Nigudahs to the community so that bad Nigudahs presumed to cause small pox can be prevented from spreading (17) (Chapter 1.4, pages 12-19; Reference 17). Interestingly, BD witnessed Jagat Ghora (JG)’s JUT-based aerosol-inoculation approach against tuberculosis (TB) disease during early years (1993–1994) of clinical practice in Sualkuchi. JG, a tantric healer and practitioner of JUT asked selected subjects with pulmonary TB (PTB) to perform Kirtan chanting in Hatisatra namghar (a local temple where the idea of KaviKrishna laboratory was initially conceived) to transmit good nigudahs in the community. JG was supported by KRD (one of the co-authors), a poet-philosopher who wrote several poems based on JUT and contributed to revive the metaphysics of JUT (25). Importantly, BD noted that most of the PTB cases managed by JG were smear-negative PTB cases. Obviously, we had direct conflict with Tantrik JG’s approach on managing PTB with JUT based approach, as we feared that such practice might spread the pathogen in the community.

However, we also considered that dialogue with IKS based traditional healers might lead to the discovery of undiscovered public knowledge (UPK) (26), which may contribute to advancing our understanding of PTB. Such dialogue may be conducted by qualitative research tool such as focused group discussion (FGD), an useful tool for health care practitioner to enhance care for patients and community (27, 28).We have been applying this FGD based research tool in conservation research (29), as well as clinical care for cancer patients in rural area of Assam (13). We also speculated that guided-interaction between IKS and modern medical practice may lead to the emergence of novel insight about disease pathology (30). Hence, we sought to utilize FGD to conceptualize good-Nigudah of aerosol-inoculation, and then apply modern laboratory-based methods to study the potential rationale for aerosol-inoculation.

Thus, FGD led us to conduct a contact investigation study among JG’s aerosol inoculation subjects, and the eventual identification of ESAT-6 rich extracellular vesicles in the aerosols of smear negative PTB (SN-PTB) subjects. Importantly, upon intranasal injection of aerosol-EVs of SN-PTB subjects, recipient mice developed immunity against a laboratory strain of *Mtb*. Moreover, broncho-alveolar lavage (BAL) fluid of these immunized mice exhibited the presence of *Mtb* infected CD45 negative cells having “altruistic stem cell” (ASC) phenotype, and capable of secreting ESAT-6 rich EVs. These mice actively secreted ESAT-6 rich EVs in their aerosols in an ASC dependent manner, as the reduction of these cells by a small molecular inhibitor FM19G11 led to decrease of EV rich aerosol production. These data indicate the potential role of a newly identified innate ASC defense mechanism (31) in the export of antigen-rich EVs by aerosol for possible spread of herd immunity. We then demonstrated that viable but not culturable (VBNC) deep dormant *Mtb* isolated from smear-negative subjects could reprogram human MSCs to ESAT-6 EV secreting ASCs. Finally, western blot approached confirmed the presence of CD63 rich ESAT-6+ rich EVs mainly in the SN-PTB subjects who’s sputum contain VBNC. In this manner, we were able to identify a group of SN-PTB subjects as the potential agents of JG’s tantric healing practice to spread good-nigudah or ESAT-6+ EVs via aerosol-inoculation. This discovery may enable to develop vaccine candidates by further studying the innate ASC based defense mechanism and their putative role in herd immunity.

## RESULTS

### A 15 years of contact TB investigation study suggests relevance of a tantric metaphysical concept of herd immunity-the Nigudah-immunity

In ancient civilization-culture based nation states such as India and China, many centuries old Indigenous knowledge system (IKS) continue to thrive as undisclosed public knowledge (UPK) of public health importance (14, 30). Body-fluid based inoculation was practiced in ancient China and India against smallpox, which may have encouraged the idea of vaccine development (32). However, aerosol-based inoculation was also practiced against small pox in ancient Kamarupa, India’s northeastern zone bordering China, and home to tantric healing practices since ancient time (17) (Chapter 5.6, pages 222-231, Reference 17). Re-discovering the metaphysics of such healing practices may enable us to develop innovative scientific hypothesis generation and experimental approaches to resolve challenges in medical science (30) (17) (Chapter 4.7, pages 162-168, Reference 17).

While treating an index PTB case in Sualkuchi in 1994 at KaviKrishna Clinic (the former name of KaviKrishna Telemedicine Care), we were confronted with co-author JG’s aerosol-inoculation practice, where selected cases of PTB were asked to attend Namghar (community temple for Kirtan chanting) for chanting Kirtan to spread good-Nigudah, subtle microscopic living being capable of doing upakar (altruism) to human (17) (Chapter 1.4, pages 11-19, Reference 17). Nigudah or invisible krimis are mentioned in sacred books of Atharvaveda, as well as Caraka Samhita, a 600 BC book on Ayurveda. However, we were alarmed by JG’s practice of aerosol inoculation, as it may encourage *Mtb* transmission in the community. At the same time, we were attracted to know more about the metaphysics that underlies JG’s traditional healing practice known as Jiva Upakara Tantra (JUT). Hence, we initiated a focus group discussion (FGD) with JG, poet-philosopher KRD (an expert in the metaphysics of JUT), and five other participants; focused and open-ended inductive interview and discussion were performed to gather data, and then analyzed by grounded theory (33).

As depicted in Figure 1A, we were able to formulate the metaphysics behind JG’s good nigudah based aerosol inoculation against PTB. The epistemology of this metaphysics, the “satvata tarka” or dialectic method of reasoning is described in the supplementary note 1. The metaphysics is based on an Ayurvedic concept of “Vyadhikshamatwa” (equivalent to immune defense mechanism) (34–36) that body defense mechanism of a TB subject can extract good-nigudah (mentioned in ancient Hindu and Jain sacred texts) from bad-nigudah (pathogen itself as per JUT-UPK) of PTB subjects and spread these extracted good-nigudha via aerosol inoculation during Kirtan chanting to devotees. Then, this good-nigudah provides protection to the community against bad-nigudah spread via aerosol from other PTB subjects. In JUT, the body defense mechanism of good-nigudah is known as the “Nigudah-Ojaskshayamatva pravriti-arthapati”, which can be roughly translated as the hypothesis (pravriti-arthapati) of “Nigudah-immunity”.

**Figure 1:**
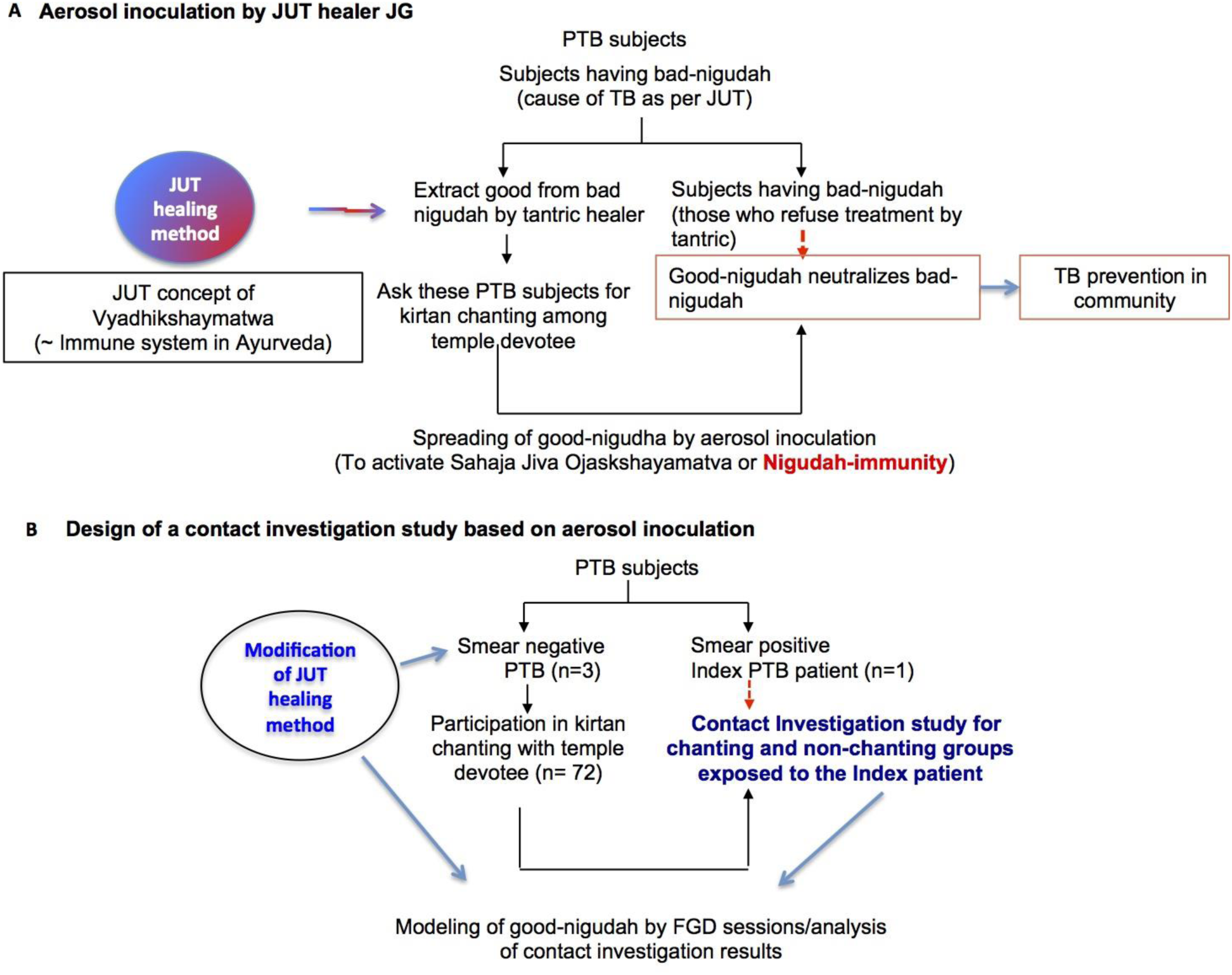
Design of a contact Investigation study to evaluate an indigenous method of aerosol inoculation against PTB. **A**: **Result of the first FGD session: discovery of Jiva Upakar Tantra (JUT) or Vedic Altruism based aerosol-inoculation method to prevent the spread of TB** (Raja yakshma in Ayurveda). The aerosol-inoculation is based on the ancient concept of Nigudah krimis (invisible krimis or microscopic life form as per Ayurveda, also known as nigoda in Jain sacred texts) as per JUT (pages 8-19 of ref. 17). This theory, known as Nigudah-Ojaskshayamatva or Nigudah-immunity is similar to Vyadhikshaymatwa or Ojaskshyamatwa, a basic concept of Ayurvedic immune defense mechanism (34; 35; 36). A JUT-based traditional medicine man (tantric ojah) such as JG extracts good-nigudah from bad-nigudah from a PTB subject. The recovered subject is then employed as a source of aerosol-inoculation to spread good-nigudah in the community (the treatment method was originally used against small pox) (pages 8-19, Reference 17). **B. Decision of the first FGD session: modification of JG’s healing practice to facilitate a contact investigation study.** Following the FGD session with JG and KRD, we modified JG’s healing practice so that he includes only the smear-negative PTB subjects in aerosol-inoculation practice; we agreed to perform a contact investigation study to find the efficacy of his aerosol-inoculation in PTB prevention among the temple devotee of Hatisatra (a 17^th^ century Hindu temple complex based on the Ek Sarana Nama Dharma of Sri Sankardevahttps://en.wikipedia.org/wiki/Sankardev).

Although, the concept of Nigudah-immunity fascinated us, we feared that such an aerosol-inoculation approach would spread TB in the community. Therefore, during the FGD session, we suggested JG to not include smear positive PTB (SP-PTB) subjects as bacteria is present in sputum of SP-PTB subjects. Further, we helped JG to modify his alternative medical practice so that he would take only SN-PTB subjects for the aerosol inoculation practice. As a trade-off, we agreed to conduct a contact TB investigation study among the Kirtan chanting devotees. JG and KRD wanted us to use the result of the contact investigation study for potential modeling of good-nigudah as per JUT’s method of “tarka” (Figure 1B, Supplementary note 1), an investigative tool of ancient India’s Nyaya school of philosophy (37). Thus, guided-interaction between IKS and modern medical practitioner led to the emergence of an experimental design of a contact TB investigation study to test the “**nigudah-immunity**” hypothesis.

Examination of close family and social contact of index cases at the time of diagnosis is a valuable method of case-findings as well as to study the disease transmission process (38). We selected an index case of smear positive PTB; a 32 year old male living in south-western part of Sualkuchi. Many people being in contact with the index case were also attending Kirtan in local Namghar (temple), Iswar Sri SriHatisatra (https://www.facebook.com/hatisatra/?rf=823873277776650), a spiritual tradition based on Sri Sankardeva’s Vaishnava tradition (107). For initial 5 years (1994–1999), we incorporated 110 subjects for contact TB tracing study among the temple devotees. Of 75 regular Kirtan chanting (once daily for average 1 hour) subjects, 3 (4%) were SN-PTB subjects (not under regular treatment) and 20 (26%) were in close contact with a SP-PTB subject. Another 35 people were found to be in close contact with the smear positive PTB subject and they were not attending Kirtan chanting. We compared the prevalence of PTB among these Kirtan-chanting (n=20) and non-Kirtan chanting groups (n=35) who came in close contact with the smear positive (SP) index case. After 5 years, 3/35 of non-Kirtan chanting versus 0/20 of Kirtan chanting group was infected with PTB (Figure 2). After 15 years (1994–2009) of follow up, the subjects who did not attend Kirtan chanting had average of 9% chance of infection with PTB following exposure to the index case, which is consistent with national data on contact-tracing (39). On the other hand, no subjects in 15 years attending regular Kirtan chanting get infected with PTB following exposure to the index case (Figure 3A & B). This result, although preliminary, and of limited use to draw a conclusion (because of small study population) encouraged us to conduct another FGD session of “tarka” to model good-nigudah, as described below.

**Figure 2:**
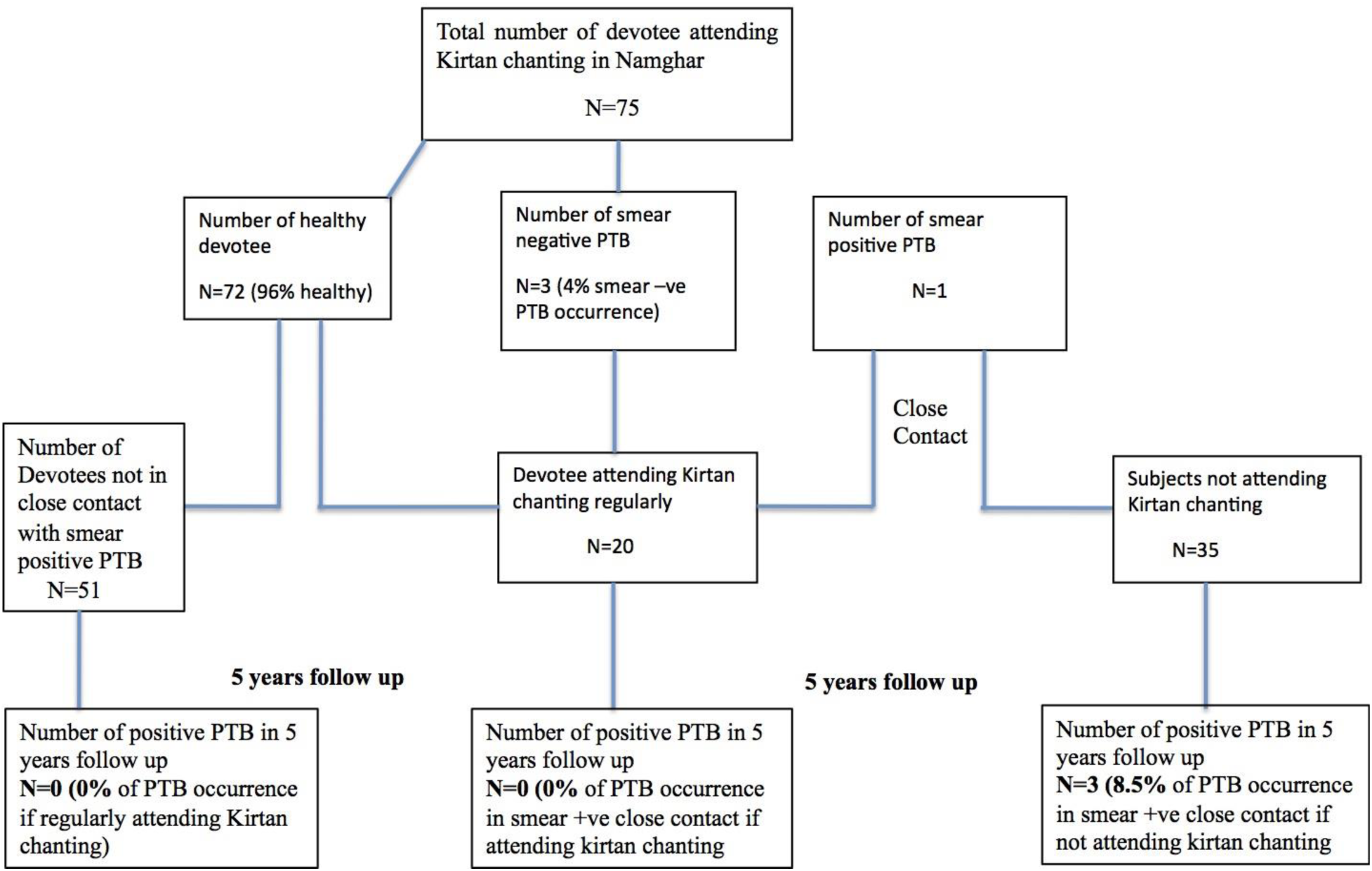
Results of contact investigation study of smear-negative PTB subjects (1994–1998). Contact investigation study was conducted among Kirtan chanting group exposed to smear negative and smear positive subjects. For initial 5 years (1994–1999), we incorporated 110 subjects for contact TB tracing study among the temple devotees at Hatisatra (Ishwar Sri Sri Hatisatra, a 17^th^ century Vaishnav temple complex in Sualkuchi (https://en.wikipedia.org/wiki/Sualkuchi), a cultural zone of ancient Vedic Kamarupa (https://en.wikipedia.org/wiki/Kamarupa) where JUT based medical practice evolved (Chapter 2 and 5, Reference 17).Of these 75 regular Kirtan chanting (once daily for average 1 hour) subjects, 3 (4%) were smear negative PTB subjects (not under regular treatment) and 20 (26%) were in close contact with a smear positive PTB subject. Another 35 people were found to be in close contact with the smear positive PTB subject and they were not attending Kirtan chanting. We compared the prevalence of PTB among these Kirtan-chanting (n=20) and non-Kirtan chanting groups (n=35) who came in close contact with the smear positive index case.

**Figure 3A.**
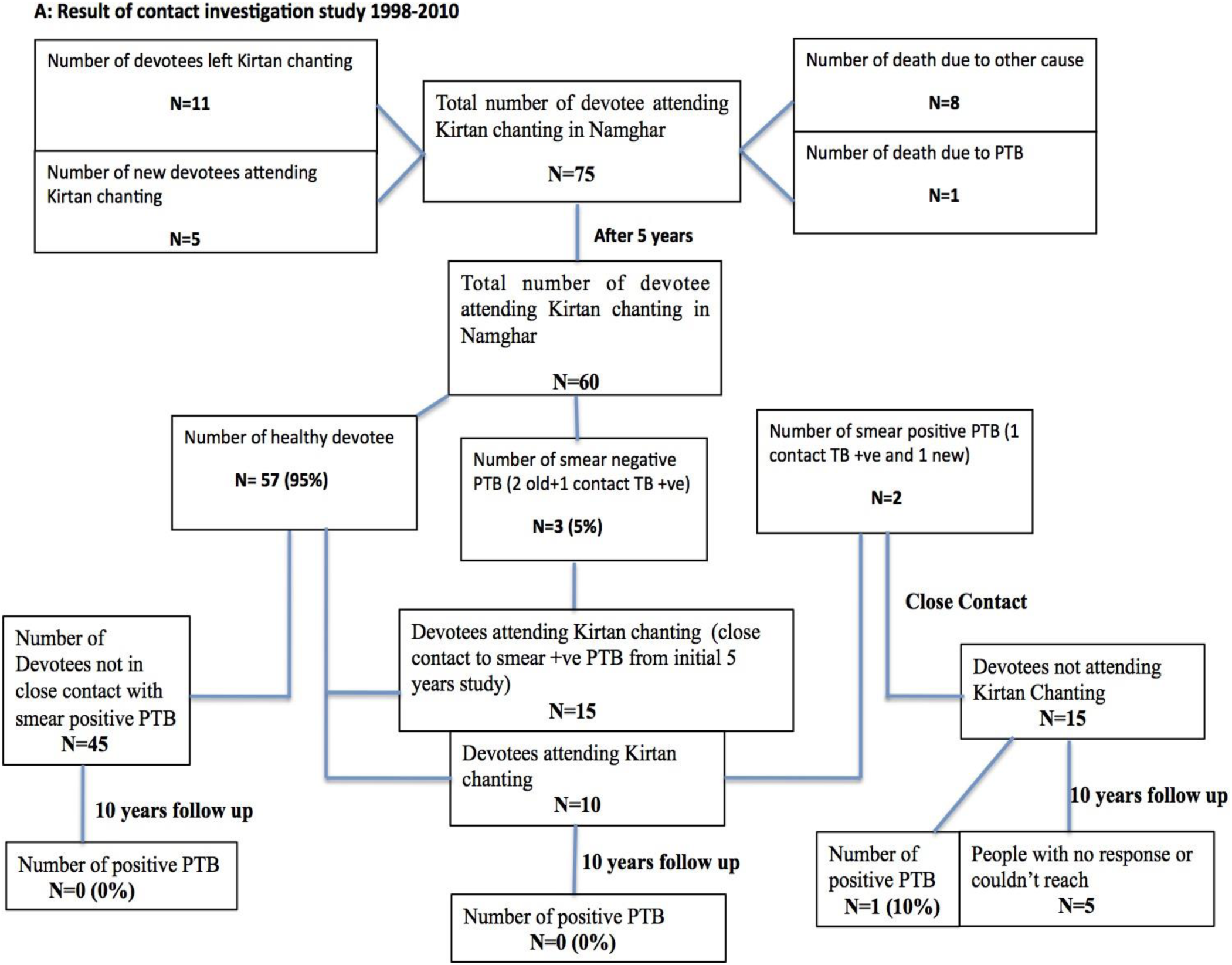
Flowchart describing the results of contact investigation study (1998-2010). After initial 5 years of contact TB investigation, 10/75 (14%) left Kirtan chanting, 8 (10%) died due to other causes and 1 (1%) died due to PTB. We initiated with 60 regular Kirtan chanting devotees for next 10 years (1999–2009) follow up, out of which 15 were the old Kirtan chanting devotees that were in contact with the smear positive PTB subject. At the end of the 10 years follow up, the 5/20 (25%) subjects left Kirtan chanting, 15/20 (75%) subjects were found to be not infected with PTB. 3/60 (5%) subjects were smear negative PTB and 10/60 (16%) were newly identified as in close contact with 2 smear positive PTB subjects. These 10 people were also not found to be infected with PTB. Another 15 people, who did not attend Kirtan-chanting, were also found to be in close contact with the 2 smear positive PTB subjects. Of 15 people, 5 (33%) were non-responder as they migrated from the village; out of the remaining 10 (66%) people, 1 (10%) person developed smear-positive PTB. A few of regular Kirtan chanting devotees died due to old age or other illness and new devotees joined Kirtan chanting in 15 years, but none of them were infected with PTB.

**Figure 3B.**
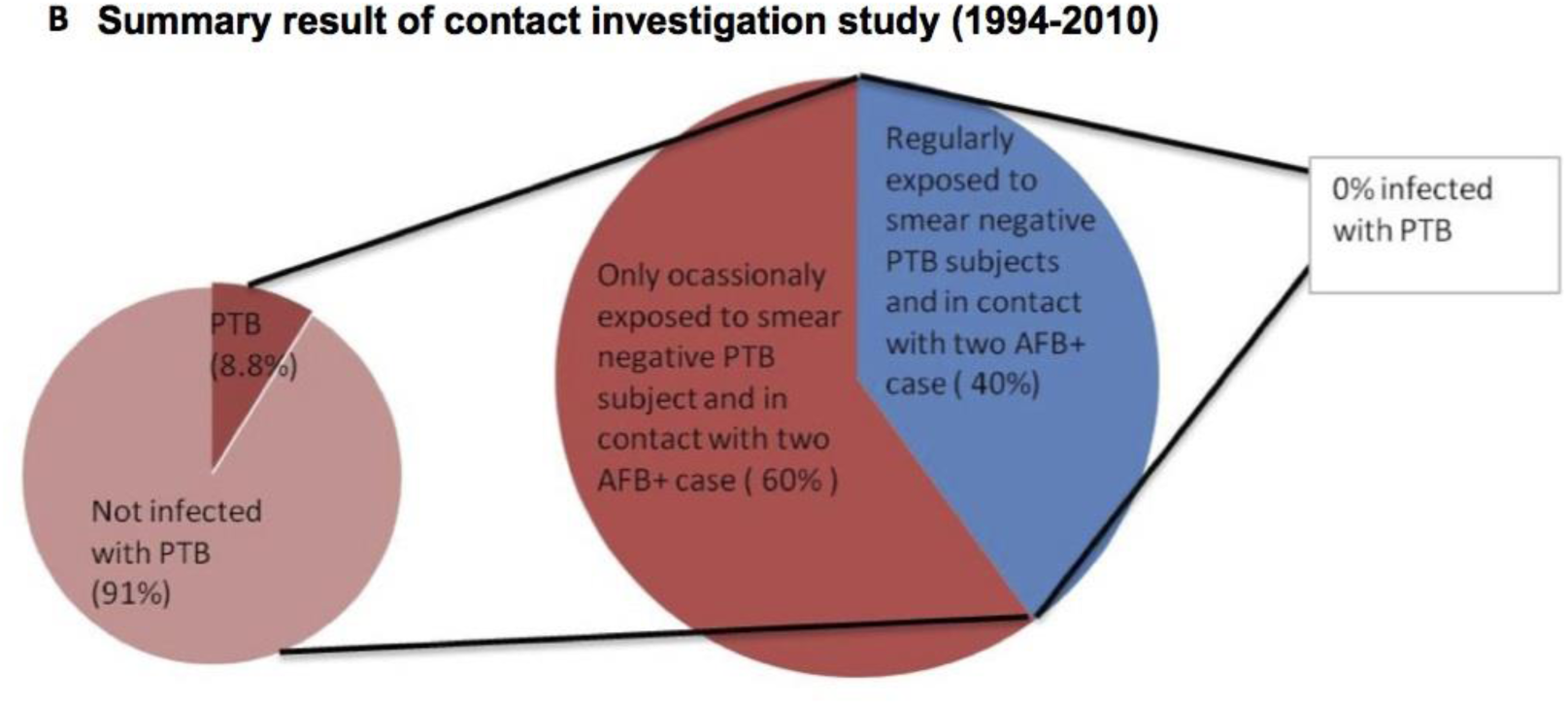
Summary pie-chart result of the contact investigation study. Blue color denotes the subjects getting aerosol inoculation while attending Kirtan chanting at the local temple. Red color represents subjects not attending Kirtan chanting or not exposed to aerosol-inoculation.

### Metaphysics based modeling of good-nigudah led to the isolation of ESAT-6 rich extracellular vesicles in the aerosols of SN-PTB subjects

Deriving a scientific hypothesis from metaphysical concepts is challenging (40) but may lead to knowledge emergence, a network process of interaction among the practitioners of local indigenous knowledge and practical philosophy (14). “Satvata tarka”, the JUT approach of interaction is a Hindu metaphysical method of reasoning and performing this method may lead to knowledge emergence may lead to knowledge emergence (17) (Chapter 2.2, pages 42-47). Thus, we performed another Satvata-tarka based FGD session to analyze the results of the 15 years of contact investigation study as per JUT tradition of Satvata-tarka (details in method section: suppositional reasoning (tarka) based modeling of good-nigudah). Because, JG, and KRD passed away, DKB, an expert on Satvata tarka participated in the FGD. The special focus of the FGD session was to model good-nigudah so that an experimental design can be formulated to identify the putative good-nigudahs from the SN-PTB subjects. Briefly, through different elements of suppositional reasoning or satvata tarka (Figure 4A, and Supplementary note 1), DKB and BD both engaged in a dialectic process to discuss two seemingly opposite interpretations of the contact-investigation study result: 1) As per BD: smear negative subjects might have spread bacteria as good nigudah to other subjects (41) through Kirtan chanting leading to spread of immunity against the pathogen (the herd immunity). 2) DKB: SN-PTB subjects might have spread good components of bacteria as good-nigudah through Kirtan-chanting leading to spread of immunity against the pathogen (Nigudah-immunity, Figure 1A). In the FGD session, as detailed in Supplementary Note 1, a metaphysical modeling approach (based on a Vedantic theory of Praibimavada, or the theory of reflection; https://en.wikipedia.org/wiki/Pratibimbavada) led us to visualize good-nigudah as extracellular vesicle (EV) rich in *Mtb*-antigen (Figure 4B. Details in method section: suppositional reasoning based modeling of good-nigudah).

**Figure 4.**
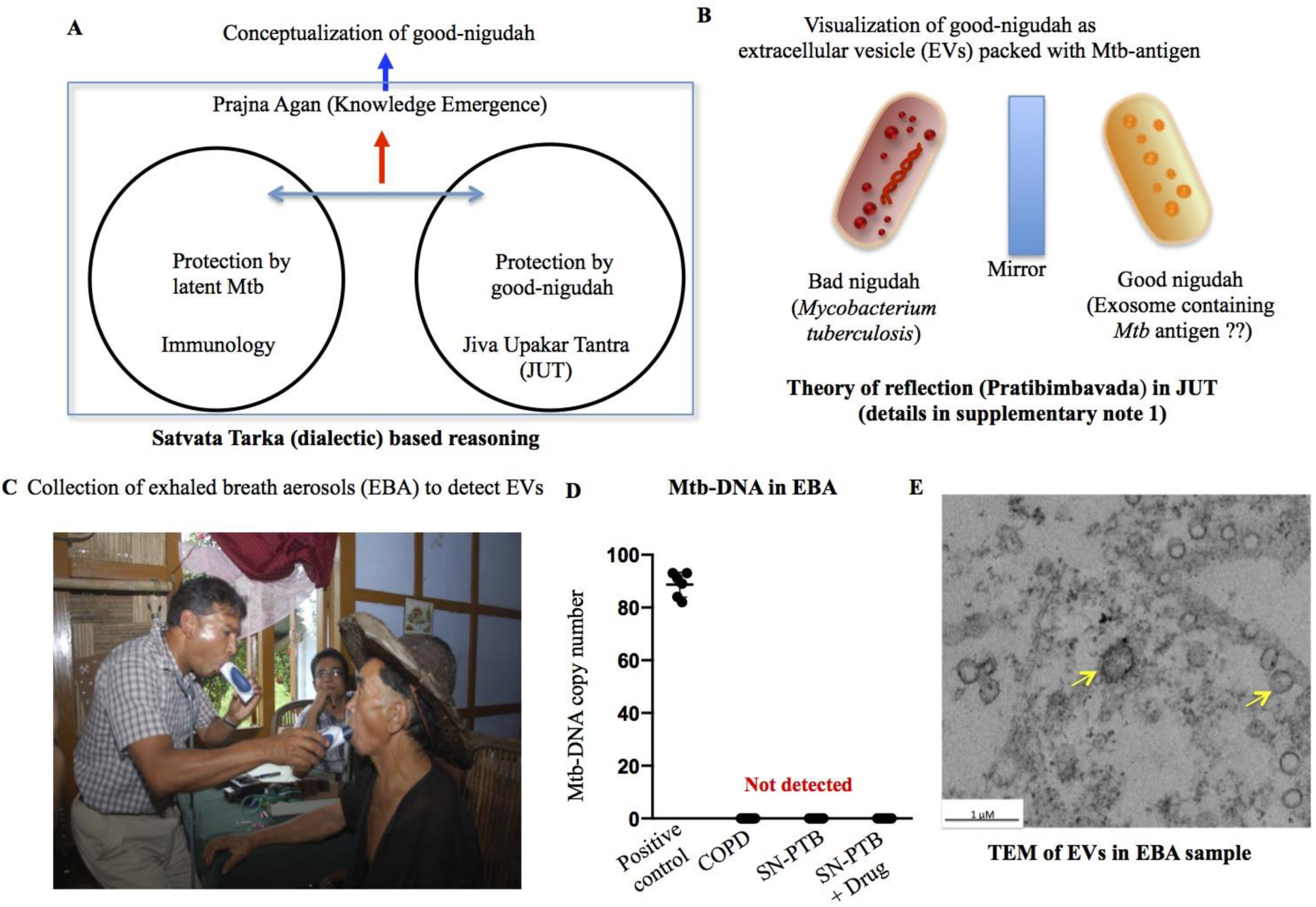
Experimental plan to identify extracellular vesicles in aerosol of smear negative PTB subjects based on the metaphysical modelling of good-nigudah. **A&B.** Results of the second FGD session. A schematic representation of the components of Satvata tarka (dialect) based reasoning. Two circles represent the first component; the structured arguments between two opposite ideas through a five-stage concentric (vyuha) process called guru-asana vyuha (the concentric seat of a guru). Thus, each circle represents seven concentric stages; in each stage, the ideas go through “parnima” (transformation) to sharpen each other’s “yukti” (reasoning). The second component is the emergence of “prajna agan” (knowledge emergence) as a result of the dynamic interaction between the two circles (Supplementary note 1). This satvata tarka was performed to debate the results of contact investigation study: chanting showed apparent protection from PTB. BD argued from the Immunology point of view (the Immunology circle) that apparent protection was due to latent infection that the subjects acquired from potential PTB subjects during chanting. This idea was opposed by DKB, who provided arguments based on Jiva Upakar Tantra (JUT): the apparent protection was due to spread of good nigudah. As shown in the top of the schematic, the tarka process led to the emergence of a metaphysical concept of good-niguda has *Mtb*-antigen rich extracellular vesicles (EVs) shown in “**B**”. In this modelling, the application of the theory of reflection (Upakar Pratibimvavada, a metaphysical view of JUT, reference 17) led to the drawing of a good-nigudah. Details are given in supplementary note 1. **C.** Photo showing the collection of exhaled breath aerosols (EBA) of smear negative PTB subjects, and **D.** EBA collection of smear negative PTB and chronic obstructive pulmonary disease (COPD) subjects are free of *Mtb*-DNA. For the positive control, 150 *Mtb*-18b were pulsed to an EBA sample (10 μg/ml protein in 100 μl PBS). *Mtb*-DNA copy number was evaluated as previously described (1). **E.** Representative resin-embedding TEM image on EVs from ultrafiltration (100kDa membrane) followed by Ultracentrifugation (110,000xg for 75 minutes at 4C; Beckman Coulter L100k) (53), smear-negative subject #2, Table 1.

Growing bodies of research suggest the important role of extracellular vesicles (EVs) in the initiation and maintenance of adaptive immunity (42, 43). EVs are bioactive vesicles formed either as microvesicles (100 nm to >1 uM), exosomes (30-150 nm) or ectosomes (100-500 nm). EVs in the body fluid or aerosols may be isolated by several methods including ultracentrifugation, density-gradient centrifugation as well as immunomagnetic sorting using EV specific marker such as CD63 (44, 45). The isolation process can be evaluated by measuring the dynamic light scattering of collected EVs, and also visualizing under transmission electron microscopy. We isolated EVs from the exhaled breath aerosols (EBA) of smear-negative PTB subjects (SN-PTB; n=10; table 1) with mild to moderate chronic obstructive lung disease (COPD) (FEV1/FEV6 <0.3, table 1), a clinical condition that we observed among the smear negative PTB subjects that participated in the contact investigation study. In COPD, EVs can be detected in the aerosols (46). In these SN-PTB subjects, exhaled breath aerosol (EBA) was collected by a microspiromter (Figure 4C) that simultaneously assessed the COPD status of these individual (FEV1/FEV; table 1). The EBA samples were devoid of *Mtb*-DNA (Figure 4D). In a similar manner, we collected EBA of COPD (n=8) and SN-PTB subjects who completed anti-TB therapy (n=8); Figure 4D and table 1. Thus, we collected EBA samples of 34 subjects; the samples were devoid of alpha amyalase (data not shown), confirming the negligible saliva contaminants of the collected EBA samples. Then, putative EVs in the EBA product were isolated by an ultracentrifugation method that we previously established to extract cell membrane to study ligand-receptor activity (47). Dynamic light scattering measurement of the EV size revealed small to medium-sized vesicles (mean diameter of 45.6 +/-7.4 nm; n= EBA of six subjects). Transmission electron microscopy revealed 40-250 nm membrane-enclosed vesicles (Figure 4E), thus confirming the EV isolation method by ultracentrifugation.

**Table 1:**
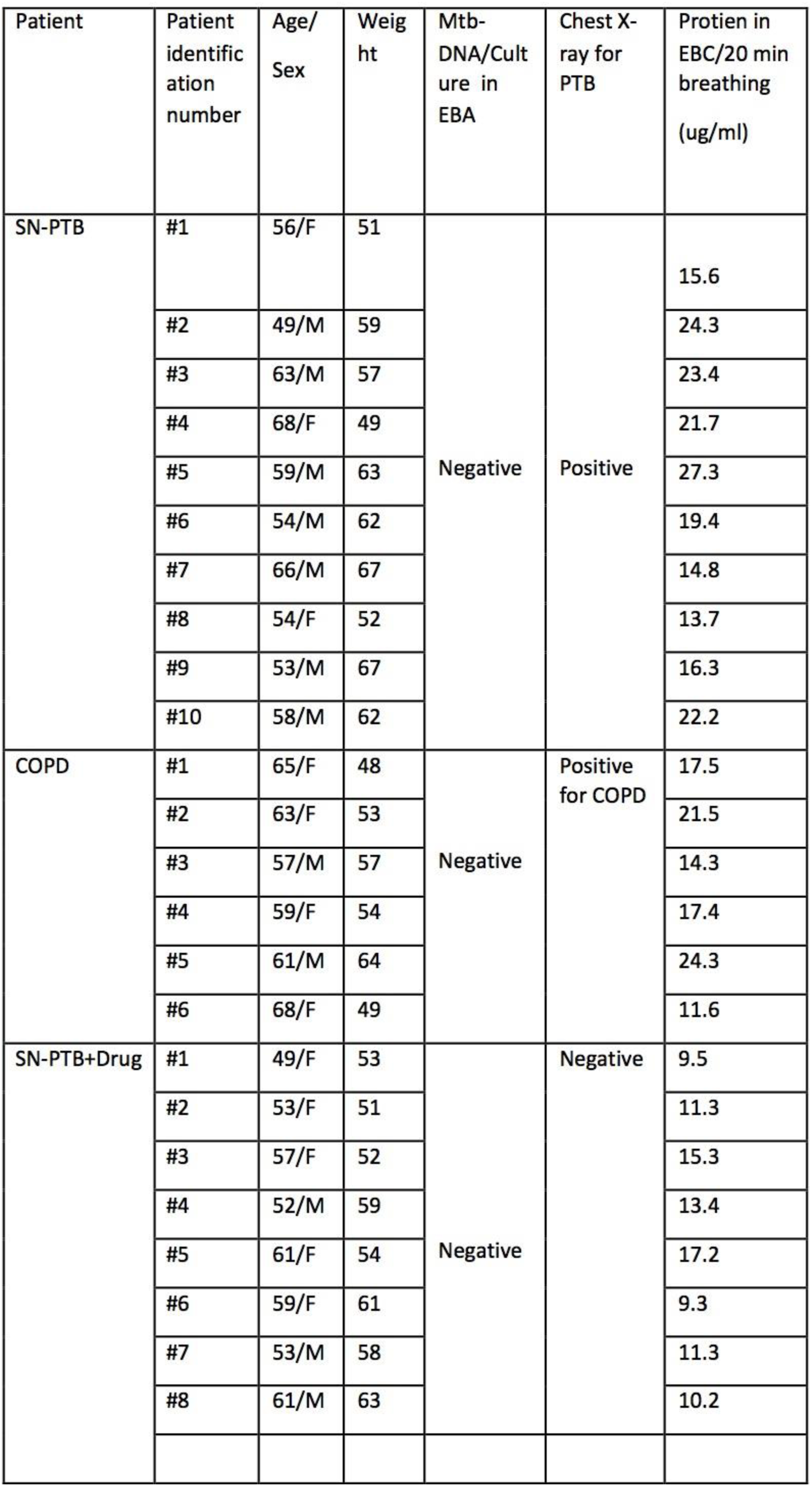
Study subjects.

| Patient | Patient identification number | Age/<br>Sex | Weight | Mtb-DNA/Culture in EBA | Chest X-ray for PTB | Protein in EBC/20 min breathing (ug/ml) |
| --- | --- | --- | --- | --- | --- | --- |
| SN-PTB | #1 | 56/F | 51 | Negative | Positive | 15.6 |
|  | #2 | 49/M | 59 |  |  | 24.3 |
|  | #3 | 63/M | 57 |  |  | 23.4 |
|  | #4 | 68/F | 49 |  |  | 21.7 |
|  | #5 | 59/M | 63 |  |  | 27.3 |
|  | #6 | 54/M | 62 |  |  | 19.4 |
|  | #7 | 66/M | 67 |  |  | 14.8 |
|  | #8 | 54/F | 52 |  |  | 13.7 |
|  | #9 | 53/M | 67 |  |  | 16.3 |
|  | #10 | 58/M | 62 |  |  | 22.2 |
| COPD | #1 | 65/F | 48 | Negative | Positive for COPD | 17.5 |
|  | #2 | 63/F | 53 |  |  | 21.5 |
|  | #3 | 57/M | 57 |  |  | 14.3 |
|  | #4 | 59/F | 54 |  |  | 17.4 |
|  | #5 | 61/M | 64 |  |  | 24.3 |
|  | #6 | 68/F | 49 |  |  | 11.6 |
| SN-PTB+Drug | #1 | 49/F | 53 | Negative | Negative | 9.5 |
|  | #2 | 53/F | 51 |  |  | 11.3 |
|  | #3 | 57/F | 52 |  |  | 15.3 |
|  | #4 | 52/M | 59 |  |  | 13.4 |
|  | #5 | 61/F | 54 |  |  | 17.2 |
|  | #6 | 59/F | 61 |  |  | 9.3 |
|  | #7 | 53/M | 58 |  |  | 11.3 |
|  | #8 | 61/M | 63 |  |  | 10.2 |

To test our hypothesis that subject with SN-PTB may export *Mtb*-antigen rich EVs in their aerosol, the isolated EVs were subjected to ELISA to detect CD63, a marker of EVs. For the candidate *Mtb*-antigen, we selected the early-secreted antigenic target 6 (ESAT-6) that has been previously detected in aerosols of PTB subjects (48). EBA of healthy adult (n=7) served as control. The results are given in Figure 5 A-B, which clearly shows that the EVs of SN-PTB subjects contain both CD63 and ESAT-6, whereas EVs of COPD subjects contain only CD63 protein. On the other hand, EVs of drug treated SN-PTB subjects showed 2-fold less amount of ESAT-6 protein compared to SN-PTB subjects (p<0.001; Figure 5B). Thus, it appears that EVs of SN-PTB subjects are enriched in ESAT-6 antigen.

**Figure 5:**
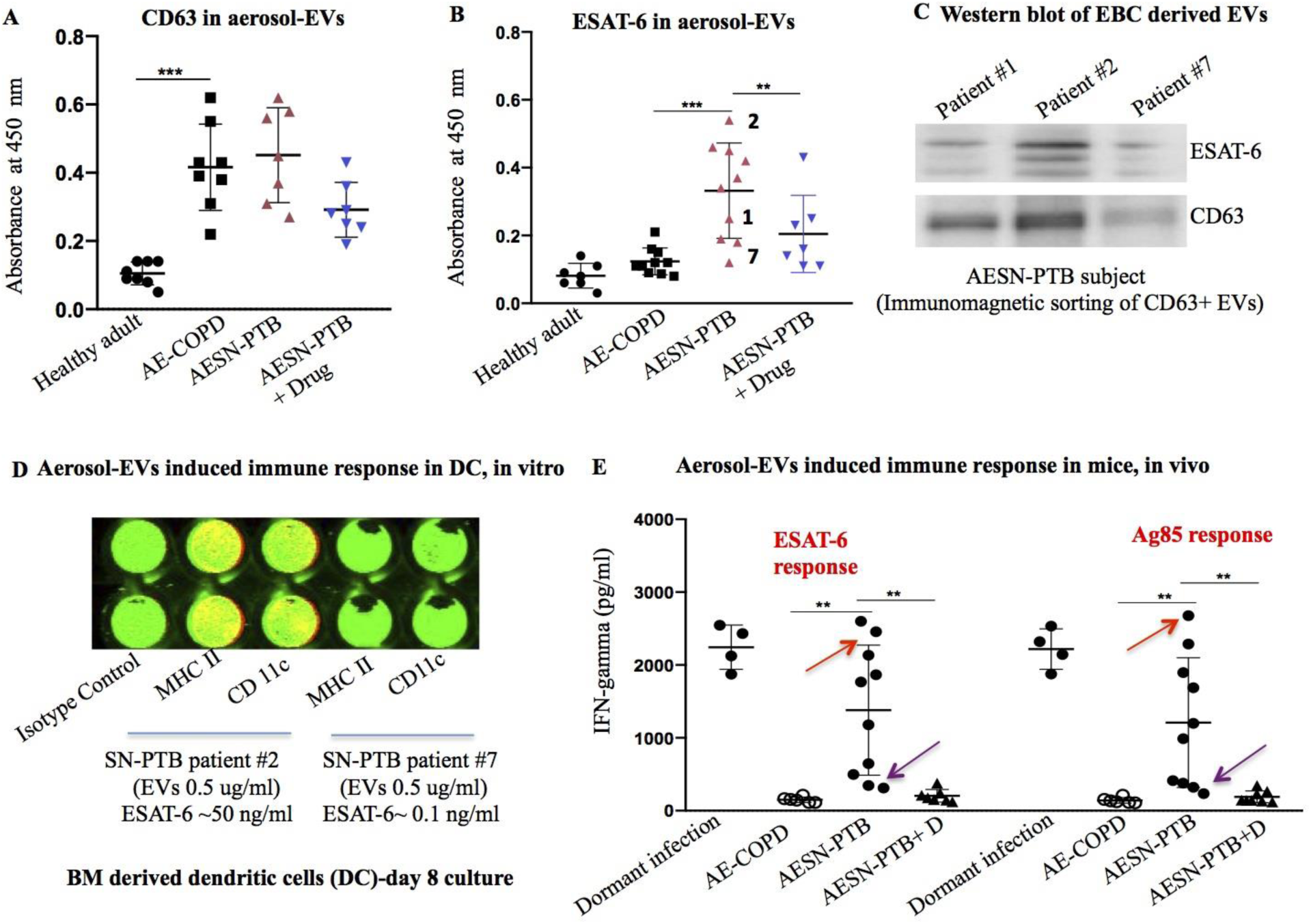
Aerosol of smear-negative PTB subjects contain ESAT-6 rich EVs capable of inducing immune response in mice. **A&B**.CD63 and ESAT-6 antigens were detected in aerosol-EVs by ELISA. For all the samples, equal amounts of EVs (8μg/ml protein) were loaded in each well. AE-COPD: Aerosol-EVs of COPD subjects; AESN-PTB: Aerosol-EVs of smear negative PTB subjects. AESN-PTB+D: Aerosol-EVs of smear negative PTB subjects completing anti-TB medications. Number 1, 2 and 7 denotes patient samples used for the western blot. **C.** Western blot of EVs obtained from smear negative PTB (SN-PTB) subjects 1-3, showing the mild, highest and lowest level of ESAT-6 protein as depicted in Figure 5B. EVs were obtained by the direct immunomagnetic sorting of CD63+ EVs in the exhaled breath aerosols (EBA), and equal amount of EVs (4 μg/ml) was loaded in each well. **D**. In Cell Western assay (53) image (merging of green and red channels), showing the induction of MHC II and CD11c expression in the mouse bone marrow derived mononuclear cells by AESN-PTB subject#2 versus #7. Equal amount of EVs (0.5 μg/ml) was added in each well, and CD11b expression (800 CW, green channel) was used for normalization of MHC II or CD11c expression (700 CW, red signal).Quantified data is given in the text. **E.** INF-γ assay in C57BL/6 mice (6-8 weeks, female). Mice were twice (a week apart) challenged with intranasal injection of 0.5 μg/ml (20 ul in 0.1 BSA) aerosol-EVs obtained from the study subjects. After 6 weeks, splenic cells were obtained and challenged with either ESAT-6 (10μg/ml) or Ag85 (10μg/ml), and then subjected to IFN-gamma measurement by ELISA. Red and chocolate color arrows represent patient#2, and 7respectively. Dormant infection was achieved by *Mtb*-m18b infection in mice (10^6 CFUs i.v. and 2 mg streptomycin/daily/two weeks) followed by two-months of streptomycin withdrawal (16960113). Data in A, B and E are presented as mean +/− SEM and each dot represents the mean value obtained from two mice injected with EVs of one study subject. **p<0.001, *** p<0.0001 (ANOVA and Dunnett post hoc test).

However, others and ours own experience suggest that ultracentrifugation method of EV isolation lead to co-precipitation of protein aggregates. Indeed, our TEM image of Figure 4E shows the potential presence of protein aggregate in the isolated EV sample. We therefore considered that ESAT-6 may also be present outside the EVs but co-precipitated during the ultracentrifugation process of EV isolation. To exclude this possibility, we obtained purified CD63+ EVs by immunomagnetic sorting technique from the 3 subjects showing heterogeneous levels of ESAT-6 protein (#1,#2,#7, Figure 5B) followed by western blotting confirmation of ESAT-6 in the sorted CD63+EVs. For this purpose, we used the exhaled breath condensate (EBC) method of aerosol collection to obtain a better yield of EVs (49) needed for the immunomagnetic based EV separation method. As shown in Figure 5C, western blot showed consistency with the ELISA results i.e. ESAT-6 content was higher in the patient +2 compared to other two patients, suggesting that the ultracentrifugation method of EV enrichment can be used for the further evaluation of EVs. We also noted that EBC versus EBA method led to 2-fold increase yield of EVs in the SN PTB subjects (4.5 ±1.3 μg/ml versus 1.8±1.1 μg/ml per 20 minutes; n=3 patients x 4 replicates, p<0.01). For further experiments, EVs were isolated by ultracentrifugation of EBC of the study subjects.

Taken together, we were able to resolve the dialect of our second FGD session with a conclusion that SN-PTB subjects might have the ability to spread good components of bacteria as good-nigudah represented as aerosol-EVs rich in *Mtb*-antigen.

### ESAT-6 + EVs of SN-PTB (AESN-PTB) exhibit potent intranasal vaccination ability in mice

For the aerosol inoculation to be effective, even a minute amount of EVs must be immunogenic and capable of activating dendritic cells present in the mucosa, and then subsequently activating the T cells in distant lymphnodes. Studies of ESAT-6 mediated intranasal vaccine (49) suggest the requirement of high doses of ESAT-6 (25 ug used in (50) to activate mucosal dendritic cells and subsequent immunization of mice. Additionally, in vitro studies shows that 5-10 ug/ml of purified ESAT-6 protein is required for the activation of bone marrow derived dendritic cells (51). Whereas, we found that in a half an hour of cumulative EBC, only ∼ 5ug of ESAT+ EVs can be collected containing about 0.1 ug ESAT-6 (Table 1). Therefore, further tests are required to find out the immunogenic relevance of EVs of SN-PTB including its potential ability to activate the Nigudah-immunity (Figure 1A and 4A).

To understand the relevance of ESAT-6 rich EVs obtained from SN-PTB subjects, we performed two experiments: first, EVs of patient #2 and #7 containing equivalent of 1-50 ng/ml ESAT-6 was added to bone marrow derived immature dendritic cells (DC) on day-6 culture (5×10^4 mouse BM mononuclear cells treated with GM-CSF 20ng/ml for 6 days; (52), and 48 hours after, In-cell western assay (53) was performed to evaluate the activation of MHC II and CD11c, two markers for DC activation. The values of MHC II and CD11c were compared with the results obtained by using recombinant ESAT-6 protein in a 50ng-10 ug dose range. While 0.5 ug/ml EVs containing 50ng ESAT-6 (patient #2) induced marked activation of MHC II and CD11c, a similar dose of recombinant ESAT-6 failed to activate DC (data not shown). Using this assay, we also noted that the similar amount of EVs obtained from different SN-PTB subjects exhibited different abilities to activate DC in vitro. A representative result is shown in Figure 5D, which shows that 0.5 µg/ml EVs containing 50ng ESAT-6 (patient #2) induced marked activation of MHC II and CD11c, whereas a similar dose of patient #7 derived EVs failed to activate DC. These results suggest the potency of selected SN-PTB subject derived aerosol-EVs (henceforth known as AESN-PTB) to induce DC activation in vitro. The findings also predict the potential ability of 0.5 µg EVs of patient #2 to induce anti-TB immunity in mice. Indeed, intranasal administration of patient #2 derived 0.5 µg EVs in 20 µl PBS weekly for two weeks showed marked increase of IFN-gamma response in mice (red arrow, Figure 5E), which is comparable to immunity exhibited by a mice model of dormant *Mtb* that we previously characterized (1). Importantly, these EV injected mice (n=3 mice per patient) showed reactivity to Ag85, a potent *Mtb* antigen being released in the aerosol of PTB subjects (54). Similar results were obtained from the patient #3-6 derived EVs (having ESAT-6 absorbance value of >0.3; Figure 5B) were able to induce immunity, whereas EVs of 4/10 SN-PTBs, as well as all the subjects of drug treated SN-PTBs failed to induce IFN-gamma response, probably because of the lack of sufficient amount of *Mtb* antigens in the EVs. Noted that the equal dose of EVs of COPD subjects did not exhibit IFN-gamma response by splenic cells, confirms that there is no non-specific IFN gamma response by EV of SN-PTB subjects.

Thus, even minute amount of aerosol EVs of some of the SN-PTB subjects were capable of generating IFN-gamma response against *Mtb* specific antigens in mice, highlighting the relevance of EV mediated antigen-export mechanism. Importantly, these findings now facilitate to develop a mouse model of nigudah-immunity as described below.

### AESN-PTB induces herd immunity in mice; towards developing a mouse model of Nigudah-immunity

The JUT’s model of Nigudah-immunity (Figure 1A/4B) provides a unique concept on herd-immunity: the export of pathogen-free antigen packed in EVs by infected individuals to the rest of a community. We have conceptualized a unique experimental model in mice that simulate the proposed mechanism of aerosol inoculation induced Nigudah-immunity (Figure 6A). Briefly, we conceptualized that in the AESN-PTB immunized mice, immune cells such as macrophages will actively produce ESAT-6 rich EVs (good-nigudah) by extracting the antigen from intracellular *Mtb* (bad-nigudah), as per the metaphysical modeling of good-nigudah (Figure 4B). These good-nigudah carrying macrophages will reach the broncho alveolar lavagae (BAL) fluid, and then into the aerosol during coughing/chanting/sneezing in human, and induced-sneezing in mice. Macrophages are the candidate host cells, as these cells are being found to secrete *Mtb*-antigen carrying EVs (55), but also the predominant cells in BAL fluid (56), a potential source for EVs in the aerosol. Additionally, MSCs, a newly identified host cells for *Mtb* intracellular dormancy and reactivation (1, 31) may also be a source of Ag-rich EVs, as these cells are well known for their EV secreting abilities (57, 58). Thus, we designed an experimental plan to look for Ag rich EVs in the BAL fluid, as well as in the aerosol of AESN-PTB immunized mice. Three weeks after immunization, these animals were infected with a laboratory strain of *Mtb* to serve as the bad-nigudah. For this purpose, *Mtb*-m18b, an *Mtb* strain that we have extensively characterized in animal models (1, 2, 31) was used. This strain can be used in BSL-2 animal facility because of its low virulence and auxotrophic nature (59). Additionally, infection with this strain can be markedly reduced by prior immunization with BCG vaccine (59). Therefore, BCG immunized mice served as a control for immunization by AESN-PTB.

**Figure 6:**
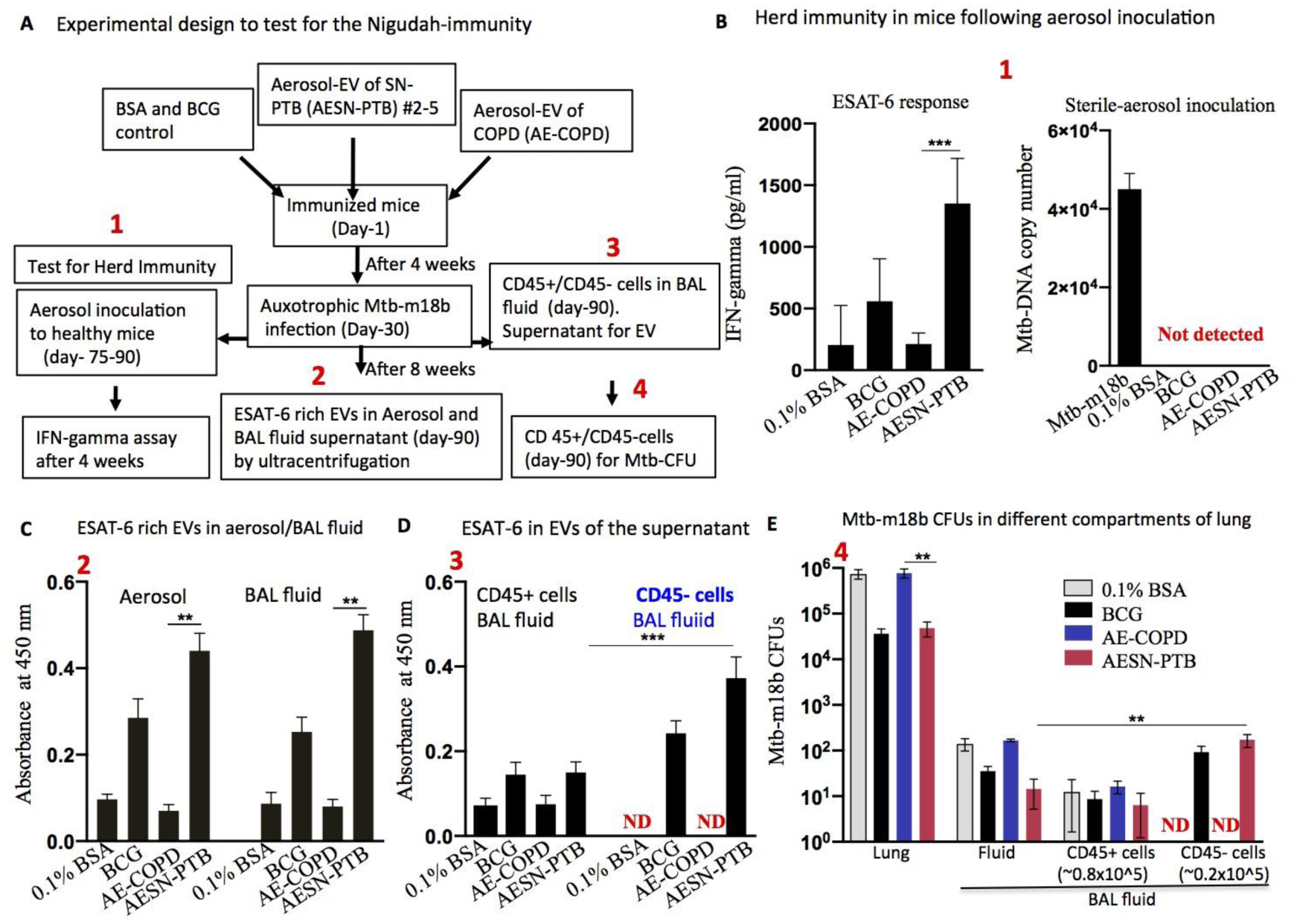
Aerosol-EVs of smear-negative PTB subjects induces an antigen-export based herd immunity (Nigudah-immunity) in mice. **A.** Schematic of the in vivo experimental plan in mice to test the hypothesis of nigudah-immunity depicted in Figure 1A-B. Mice were immunized or vaccinated with intranasal injection of 40 μl of 0.5 μg/ml/weekly/two weeks of aerosol-EVs obtained from COPD (n=15; 3 mice each for three subjects), or SN-PTB subjects(n=20; 5 for each of the four PTB subjects showing >1.5 ng/ml IFN-gamma response in Figure 5E). Control groups of mice received intransal injection of either 40 μl of 0.1% BSA (n=15 mice) or BCG (5×10^5 CFU of ATCC # 35734; n=15 mice). Red-ink numbers 1-4 represent successive data collection plan to test for vaccine-induced protective immunity against *Mtb*-m18b infection (1×10^6 CFU by intravenous injection, and daily supplemented with 2 mg of stremptomycin sulphate s.c. for 4 weeks (59, 1). **B.** Left histogram: IFN-gamma response of splenic cells obtained from healthy mice (n=4 in each group) subjected to aerosol inoculation (cumulative 50 minutes in two weeks; ∼15 μg EVs inhaled from 5 mice); four weeks after animals were sacrificed for the assay. Right histogram shows that the lungs or spleens of these mice were devoid of *Mtb*-DNA. The lungs obtained from a *Mtb*-m18b (1×10^5^ CFUs, intranasal injection and 2 mg streptomycin/two weeks) infected mice (n=3)were used as positive control. **D.** EVs isolated from the aerosols and BAL fluid supernatant of AESN-PTB vaccinated and *post-challenge Mtb*-m18b infected mice (n=5 for each of the four subjects) shows the presence of ESAT-6 protein (ELISA). **E.** CD45+ and CD45-cells of BAL fluid (Figure 6D) were in vitro cultured for 24 hours in serum free media (58), and EVs were isolated from the supernatant by ultracentrifugation to perform ELISA for ESAT-6 (total ∼2×10^5 CD45-cells were pooled from ten mice). **D.** *Mtb*-m18b CFUs of lung and BAL fluid supernatant and CD45+/CD45-cell fractions (BAL fluid of 5 mice of each subject were pooled as one, and then immunomagnetic sorting of CD45+ cells were performed as described (1). Data in B, C-E are presented as mean +/− SEM. **p<0.001, *** p<0.0001 (ANOVA and Dunnett post hoc test).

Thus, mice immunized with the AESN-PTB were infected with *Mtb*-m18b (henceforth known as nigudah mice), and 8 weeks later (a required period for infection to establish in the lung), extraction of good-nigudah (*Mtb*-Ag rich EVs) from bad-nigudah (*Mtb*-m18b) was demonstrated by several assays. First, we demonstrated that healthy mice exposed to the aerosols of these nigudah mice exhibited ESAT-6 specific IFN-gamma response without being infected with *Mtb*-m18b. We noted that neither AE-COPD nor BCG immunized mice showed the ability to induce marked IFN-gamma response in the aerosol-exposed mice (Figure 6B). Second, the nigudah mice exported ESAT-6+ EVs in the aerosols, BAL fluid, and the CD45-cells of the BAL fluid. Surprisingly, CD45+ cells in BAL fluid, enriched in macrophages did not export ESAT-6 + EVs. Third, nigudah mice exhibited immunity against *Mtb*-m18b, as the *Mtb*-CFUs in lung, and spleen (not shown) markedly decreased as compared to control groups. The number of CFUs in BAL fluid of non-immunized mice was very low in all the groups, which is consistent with previous results that *Mtb* do not disseminate to BAL fluid in C57BL/6 mice (60). BCG immunized mice also exhibited immunity against *Mtb*-m18b infection, and exhibited the export of ESAT-6+EVs by CD45-cells (Figure 6C-E), which is not surprising as BCG vaccine are known to mediate unconventional immune response against *Mtb* infection (61). The EVCOPD group of mice showed no evidence of immunity against *Mtb*-m18b, and no marked presence of CD45-cells in the BAL fluid (Figure 6 C-E). Also, AESN-PTB injected mice alone, without infection with *Mtb*-m18b, did not exhibit the presence of CD45-cells in the BAL fluid, or the secretion of ESAT-6+ EVs in their aerosols (not shown), which is expected, as injected antigen are rapidly taken by immune cells, and then processed.

Taken together, the niguda mice (animal receiving AESN-PTB intranasal injection, Figure 5E, plus *Mtb*-m18b infection, Figure 6A) exhibited a unique mechanism of herd immunity: export of ESAT-6 rich EVs in their aerosols for immunizing neighboring mice. The source of the ESAT-6 in the EVs is most likely the *Mtb*-18b, because, AESN-PTB injection alone did not cause the export of ESAT-6+ EVs (not shown). These results make this mouse model a putative model to study the proposed Nigudah-immunity.

### Nigudah-immunity recruits the innate altruistic stem cell defense mechanism

The detection of CD45-cells in the BAL fluid of nigudah mice, and their potential role in extracting good-nigudah (Ag rich EVs) from bad-nigudah (*Mtb*) made us to consider the potential role of newly identified innate ASC defense mechanism in the Nigudah-immunity. Our ongoing work indicates that CD45-cells, which are mainly enriched in MSCs may exhibit an altruistic stem cell (ASC) defense mechanism against pathogen such as MHV-1 virus. Briefly, ASCs exhibit an enhanced stemness phenotype as well as an altered p53/MDM2 oscillation to acquire a transient state of cytoprotective activity as a part of the innate stem cell defense mechanism for niche protection (2, 31, 53). ASCs are dependent on hypoxia-induced transcription factors (53) which are involved in the exosome production in MSCs (62). Notably, ASC state facilitates *Mtb* replication, and therefore, could be potential source of Ag rich EVs. Importantly, ASCs may be part of an unconventional immune response (63) against *Mtb* infection, as seen in BCG immunized mice challenged with *Mtb* infection (61, 64). Therefore, we considered to study whether Nigudah-immunity recruits the ASC defense mechanism for antigen-exports via aerosol.

To directly visualize *Mtb* harboring CD45-cells, and study their ASC phenotype in the Nigudah immunity model of mice, we used a GFP labeled *Mtb*-m18b strain (1) and recovered CD45-cells from the BAL fluid (8 weeks after *Mtb*-infection, as shown in Figure 6A) for in vitro culture, including the status of intracellular *Mtb*-18b, and the putative ASC phenotype of the CD45-cells. A group of mice were treated with FM19G11, an inhibitor of ASC phenotype (53), and the animals were evaluated for the release of ESAT-6+ EVs in their aerosols and BAL fluid. In this manner, we were able to directly visualize the *Mtb*-m18b intracellular to CD45-cells in the BAL fluid (Figure 7A). Notably, the CD45-cells appeared to exhibit a phenotyhpe of “enhanced stemness” (Figure 7A) that we recently showed to be important for dormant *Mtb* reactivation (31). Next, the intracellular bacteria number was higher than CFU counts suggesting the presence of VBNC (Figure 7B), which we confirmed by addition of early stationary phase supernatant (ESPSN) of *Mtb*-H37Ra, a conditioned media that can resuscitate viable but non culturable *Mtb* (65). Then the real time PCR based gene expression data showed the ASC phenotype of CD45-cells, including the high expression of HIFs (Figure 7C-D), which are transcription factors involved in the enhanced stemness reprograming (53). Importantly, FM19G11 treatment led to marked reduction of the CD45-ASC number, and *Mtb*-DNA, differentiation, as well as non-detection of ESAT-6 rich EVs in the cells, as well as BAL fluid and aerosols (Figure 7E). These results indicate that the putative CD45-ASCs of BAL fluid may be the source of ESAT-6+ EVs. Importantly, our findings indicate the potential role of innate ASC defense mechanism in secreting ESAT-6+ EVs. Moreover the results indicate VBNC as the potential source of the Ag in EVs, as these CD45-ASCs contained VBNC.

**Figure 7:**
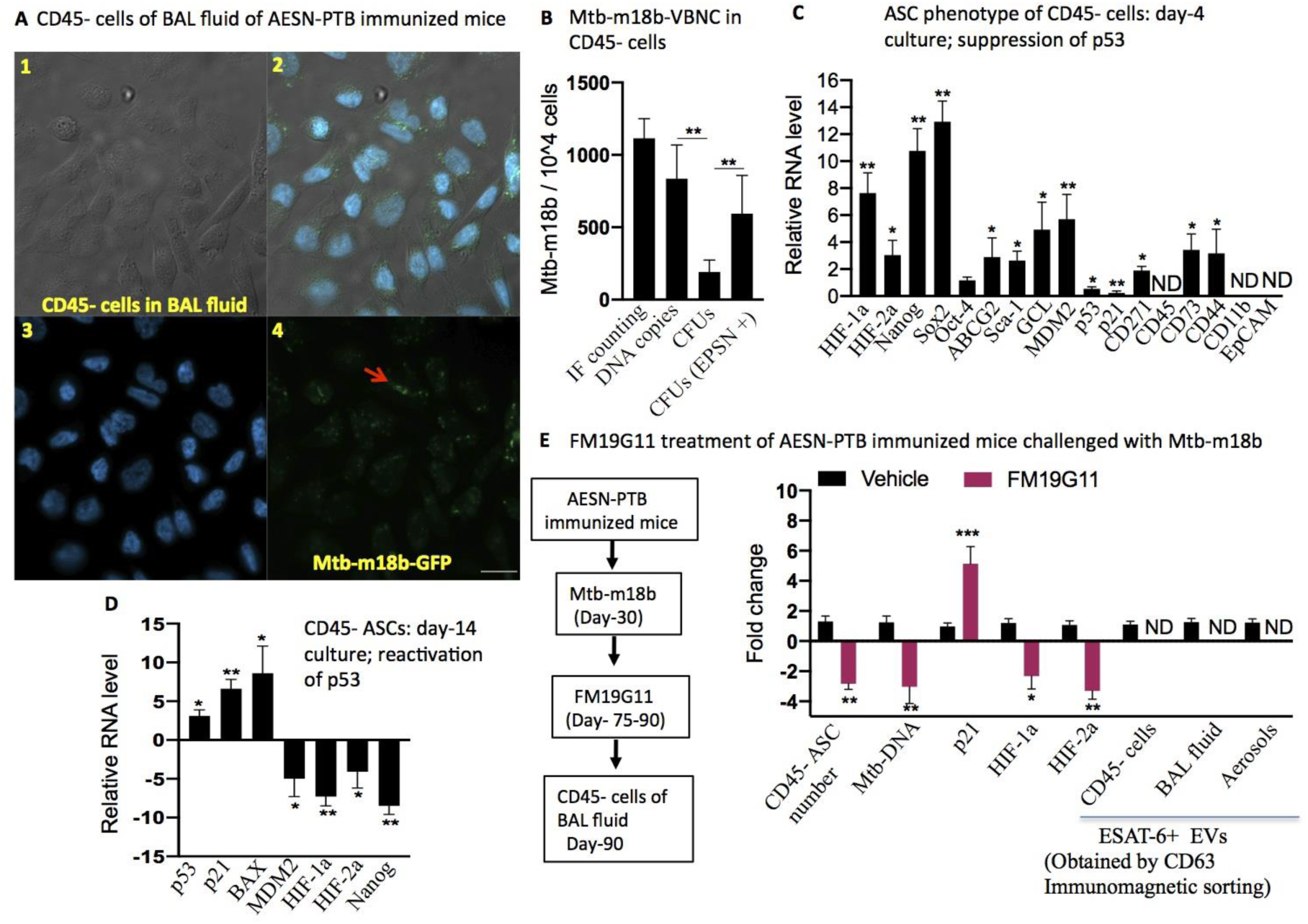
In the aerosol-EV immunized mice challenged with *Mtb* infection (nigudah-mice), CD45 negative cells of BAL fluid exhibit innate altruistic stem cell (ASC) defense by exporting ESAT-6 rich EVs. Sorted CD45-cells of BAL fluid (AESN-PTB subject #2; on day-90 after aerosol-EV immunization, and infected with *Mtb*-m18b-GFP) were either cultured on a glass chamber coated with polylysine and fibronectin for IF imaging, or cultured in 48 well plates (1×10^3 cells/well in serum free media for naïve MSCs) for functional assays. **A.** Representative phase contrast and immunofluorescence images of immunomagnetically sorted CD45-cells showing intracellular *Mtb*-m18b-GFP (red arrow). 1. Phase contrast microscopy. 2-4: IF images, merged, DAPI staining for nucleus, and *Mtb*-m18b-GFP. Scalebar 20 μm. **B.** The CD45-cell intracellular *Mtb*-m18b-GFP bacteria were manually counted per microscopic field, and the data was used to find number of bacteria per 1×10^4 cells. The data was compared with the *Mtb*-DNA copy number, and *Mtb*-CFUs with and without early stationary phase supernatant (ESPSN) of *Mtb*-H37Ra, a conditioned media that can resuscitate viable but non culturable *Mtb* (65). **C &D** Sorted CD45-cells exhibit self-sufficiency and ASC phenotype on day-8 (31) and differentiate/activate p53 on day-14, confirming the transient state of their enhanced stemness state (53;12;31). The gene expression data was compared with the MSCs of dormant *Mtb*-m18b infected mice (1). ND-not detected. **E.** Schematic of experimental design. **A**nimals (n=15; 5 mice for each subject #2-4) were injected with FM19G11 (5 mg/kg/ dissolved in 1.5% DMSO, i.p. thrice weekly for two weeks) for two weeks (12),6 weeks after the infection with *Mtb*-m18b. The control groups were injected with 200 μl 1.5% DMSO in saline, i.p. thrice weekly for two weeks. Results showed that FM19G11 treatment lead to marp>ked reduction of Nigudah-immunity, including the reduction in ASCs (CD45-cells) in the BAL fluid, and corresponding reduction in the ESAT-6+ EVs in the aerosols (key component of the Nigudah immunity). EVs (average 0.5 ug/10^5 cells) were isolated by ultracentrifugation method; Data in B,C-E are presented as mean +/− SEM. **p<0.001, *** p<0.0001 (ANOVA and Dunnett post hoc test).

Identification of ASCs harboring *Mtb*-m18b derived VBNC as the source of ESAT-6 in the nigudah-mice, led us to review current knowledge on VBNC. Although VBNC phenotype has been identified as deep dormant *Mtb* (66), lack of a good in vitro model of VBNC limit our ability to study the host/pathogen interaction between these dormant *Mtb* and host cells such as MSCs. It is important to develop in vitro model of VBNC to study if these bacteria may reprogram MSC to ASC and in the process, releases ESAT-6+ EVs.

Hence, we first developed an in vitro model of VBNC by exposing non-replicating *Mtb* (NRP II) with hypoxia/oxidative stress (67), and then used the method to obtain VBNC from *Mtb*-m18b. The VBNC of m18b was used to study the potential reprogramming of MSCs to ASC by VBNC, and associated secretion of ESAT-6+ EVs.

Thus, human CD271+ BM-MSCs were infected with *Mtb*-S18b derived VBNC, and then we studied the ASC reprogramming by evaluating p53/MDM2 oscillation within a period of two weeks, and also the expression of stemness genes. The data is shown in Figure 8A-B, which clearly showed the VBNC-induced reprogramming of MSCs to ASC phenotype. Next, we found that VBNC but not actively growing m18b infected CD271+ BM-MSCs were capable of secreting ESAT-6+ EVs, without secreting live bacteria. Moreover, inhibition of HIFs by FM19G11 (12) led to marked reduction of ESAT+EV secretion by the VBNC infected MSCs (Figure 8C-E). Thus, VBNC but not actively growing *Mtb* may acts as a source for ESAT-6+ EVs in MSCs, which needs further investigations.

**Figure 8:**
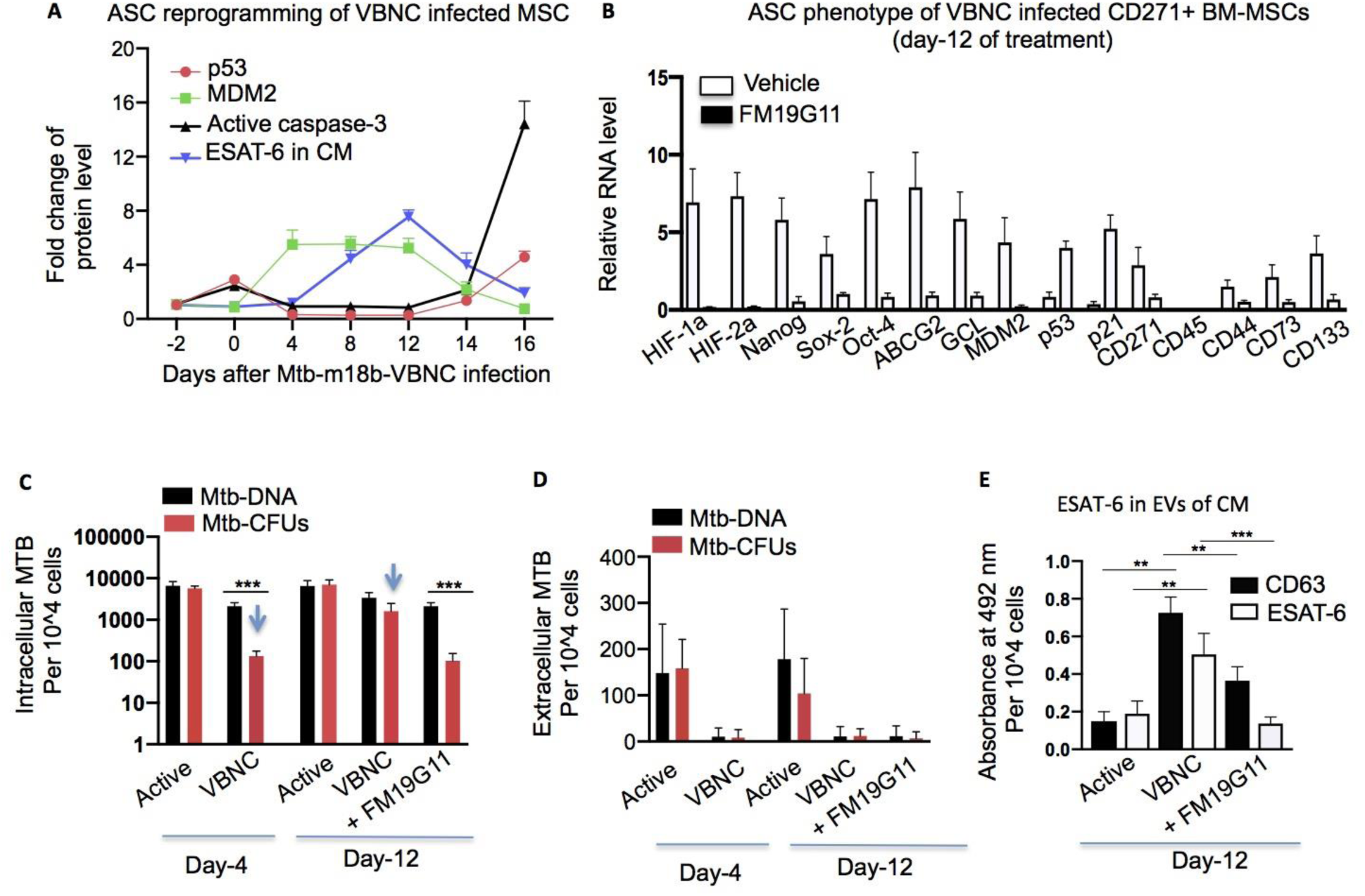
In vitro assay to study the potential role of altruistic stem cell (ASC) based defense in Nigudah immunity. **A&B: Naïve CD271+ BM-**MSCs treated with *Mtb*-m18b derived VBNC (1×10^4 /2×10^6 BM-MSCs in 6 well plates) led to enhanced stemness reprogramming, which can be inhibited by FM19G11 (0.1 μM/72 hours starting on day-9), a HIFs inhibitor known to inhibit enhanced stemness or altruistic stemness reprogramming (53). “**A**” shows an altered state of p53/MDM2 oscillation as measured by In-Cell ELISA, whereas “**B**” shows the expression of enhanced stemness genes, two key features of ASC phenotype (53; 58, 31). RNA data was compared with freshly obtained CD271+/CD45-BM-MSCs of one healthy donor. Noted that “-2” in “A” denotes cell seeding two days before VBNC addition.**C-E.**BM-MSC culture and the conditioned media of day-0-12 as shown in “A” were subjected *Mtb*-DNA, *Mtb*-CFU as well as EV isolation (by density gradient centrifugation) assays to study relation between VBNC resuscitation and the secretion of ESAT-6 rich EVs. “C” shows resuscitation of *Mtb*-m18b derived VBNC by CD271+ BM-MSCs on day-12 versus day-4 culture (blue arrows). “D” shows that the process of VBNC resuscitation did not cause extracellular secretion of *Mtb*, but (E) triggered the secretion of ESAT+ and CD63+ EVs. FM19G11 (0.1μM/72 hours starting on day-5) treated cells failed to resuscitate VBNC as well as secretion of ESAT-6 rich EVs, which was obtained by density gradient centrifugation of conditioned media of cumulative 5×10^6BM-MSCs; experiment was repeated four times. CD271+ BM-MSCs were obtained by direct flow cytometry sorting of CD271+/CD45-cells from the BM mononuclear cells of healthy donors following proper consent (1). **P< 0.001, ***P<0.0001, Student’s t-test; Error bar represents SEM.

### Smear negative PTB subjects with viable but non culturable (VBNC) *Mtb* may spread Nigudah-immunity

After elucidating VBNC-*Mtb* as the source of ESAT-6+ EVs, we revisited our clinical data of Figure 5, where the aerosols of 6/10 pretreated SN-PTB subjects exhibited ESAT-6+ EVs of greater than 0.3 absorbance at 450 nm, and also showed ability to induce the Nigudah-immunity in mice (Figure 6). We speculated that these subjects might contain VBNC-*Mtb* in their BAL fluid, and therefore in the sputum. Indeed, whereas processed sputum of these subjects were negative for *Mtb*-DNA or culture, addition of ESPSN media in the processes sputum led to actively growing *Mtb* in 6/10 samples (Figure 9A). Thus, it appears that we have identified a group of SN-PTB subjects having VBNC-*Mtb* in sputum that spread aerosol EVs containing ESAT-6 into the community. The SN-PTB post treatment subjects without VBNC-*Mtb* (n=8) served as control. As a positive control for western blot, we used the EVs released by VBNC-*Mtb* containing ASCs (Figure 8), which clearly showed the presence of ESAT-6 as well as CD63 proteins (Figure 9B). Thus, AESN-PTB having VBNC-*Mtb* in the sputum contain ESAT-6+ EVs in their aerosols. Importantly, these EVs injected nigudah-mice (Figure 6A) showed 15-fold higher number of ASCs in the BAL fluid than the mice injected with the AESN-PTB having no VBNC-*Mtb* in their sputum (p<0.0001, Figure 9C). Moreover, in these nigudah-mice, we observed a direct correlation between the number of ASCs in BAL fluid, and the amount of ESAT-6+ in the aerosol-EVs (p<0.019, Figure 9C). These results indicate that VBNC versus non-VBNC group of AESN-PTB is capable of inducing a higher degree of innate ASC defense in the nigudah mice. Our results also show that the innate ASC defense is directly correlated with the Nigudah-immunity i.e. higher release of Ag rich EVs in the aerosols of these mice for exerting herd immunity.

**Figure 9:**
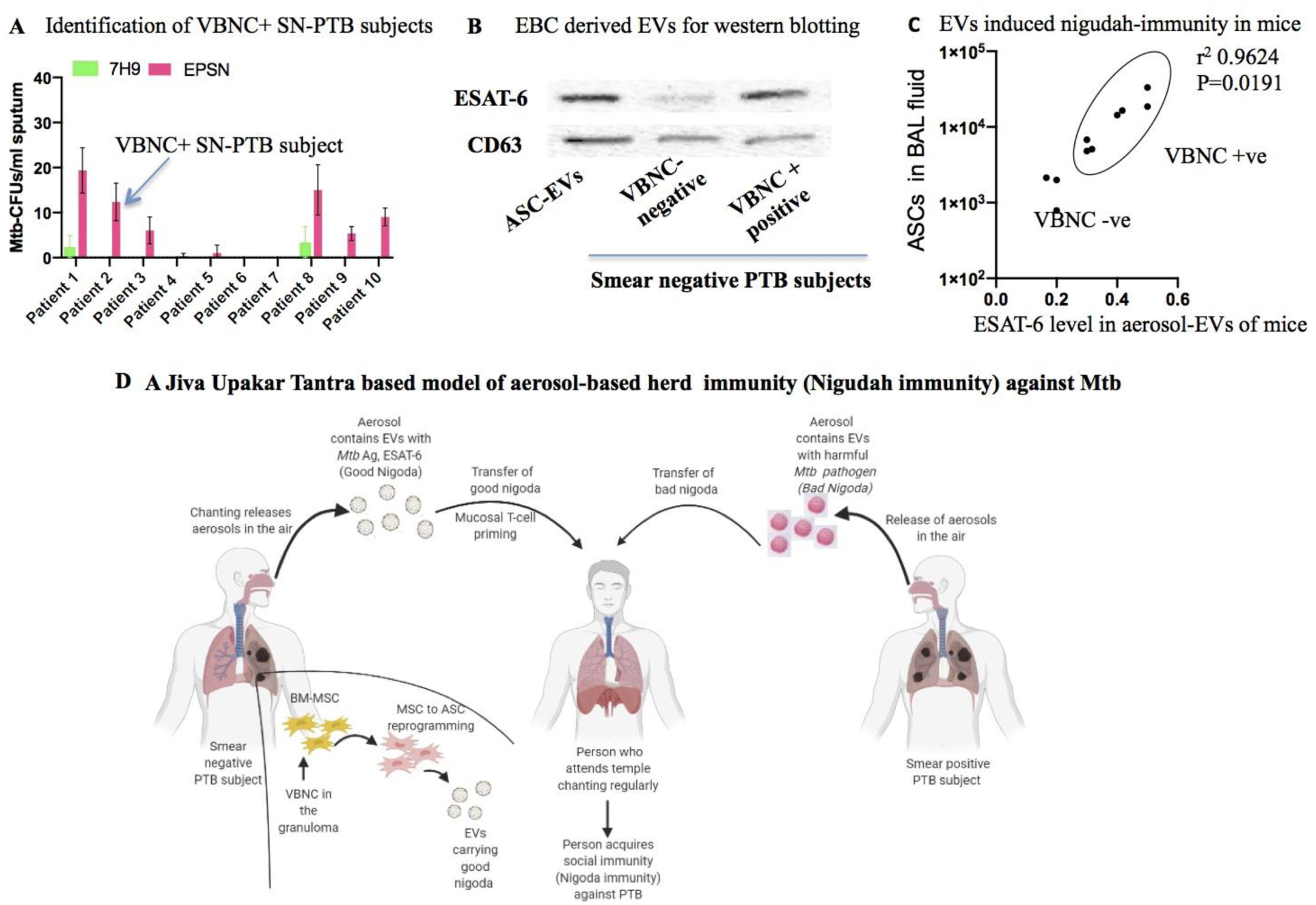
Viable but not culturable (VBNC) Mtb positive fraction of smear-negative subjects export ESAT-6 rich EVs in their aerosols. **A.** Identification of VBNC + smear negative PTB subjects by subjecting processed sputum to early stationary phase supernatant (ESPSN). The ESPSN is rich in resuscitation factors, and was obtained from *Mtb*-H37Ra as previously described (65). Smear negative sputum that exhibited *Mtb*-CFUs (as confirmed by AFB staining) following ESPSN treatment was identified as sputum containing VBNC.**B.** Western blot images of CD63 and ESAT-6 proteins present in the EVs (10 µg loaded in each well) obtained from the exhaled breath condensate (EBC) of VBNC versus non-VBNC subjects. EBC products were pooled together from the 4/6 VBNC and 4/4 non-VBNC subjects, and EVs were isolated by density gradient centrifugation method. EVs (10 μg loading in each well) obtained from CD271+ ASCs infected with VBNC (Figure 8E) served as the positive control. **C**. Correlation between the number of CD45-ASCs cells in the BAL fluid, and the optical density reading of the ELISA assay for ESAT-6 in the aerosol-EVs (5 μg/ml) of the corresponding mice. Data is obtained from the experimental results of Figure 6C&D. **D**. A schematic model of Nigudah-immunity: VBNC present in the granuloma of PTB subjects are recruited by BM-MSCs migrating to lung (108). VBNC harboring BM-MSCs reprogram to altruistic stem cells (ASCs) and migrate to the BAL fluid, where these ASCs releases EVs pack with *Mtb*-antigen including ESAT-6. Following coughing or Kirtan-chanting, these EVs spread in the community by aerosol based transmission, and come in contact with the mucosal immune cells (M cells and DC of nasal, laryngeal and bronchial mucosa) of healthy subjects, or subjects affected by active PTB. EVs efficiently deliver the *Mtb* antigen to DC and other antigen-presenting cells leading to T-cell priming. In the subjects with active PTB, EV-induced immunity may also activate the innate ASC defense mechanism (31) including the extraction of *Mtb*-antigen from the VBNC, and export antigen rich EVs into the BAL fluid.

## Discussion

Pulmonary tuberculosis (PTB) is not yet eradicated from the globe despite decades of TB control program (68). *Mtb*, the causative agent of PTB enters into the human host via aerosol and causes wide-spread infection in the community. In this context, while in a densely populated country like India half of the community is infected with *Mtb*, yet only in 5-10% infected people develops PTB (69). It is not yet clear, how the 90% of population escape from PTB. In this manuscript, we reveal an IKS based approach of aerosol inoculation in the community to prevent TB infection, and its associated metaphysics known as Nigudah-immunity (Figure 1A). We carefully integrated this healing metaphysics to design innovative experimental plan, and found evidence for a novel mechanism of natural vaccination against TB. In this mechanism, the “Nigudah immunity” as described in Figure 9D, smear negative PTB subjects export *Mtb*-antigen rich sterile EVs to the community via aerosol and these EVs are capable of exerting *Mtb* specific immunity in a mouse model of *Mtb* infection. Importantly, we uncovered the role of newly identified innate altruistic stem cell (ASC) defense mechanism (31) in the generation of Ag-rich EVs. Based on the findings, we propose that VBNC-*Mtb* present in the granuloma of PTB subjects is recruited by BM-MSCs migrating to lung (108). VBNC-*Mtb* harboring BM-MSCs reprogram to altruistic stem cells (ASCs) and migrate to the BAL fluid, where these ASCs releases EVs pack with *Mtb*-antigen including ESAT-6. Following coughing or Kirtan-chanting, these EVs spread in the community by aerosol based transmission, and come in contact with the mucosal immune cells of healthy subjects, leading to natural vaccination (Figure 9D).

Our study opens up new vista on *Mtb*/Stem cell host/pathogen interaction. Following our initial report on *Mtb* infection of mesenchymal stem cells (MSCs) as well as hematopoietic stem cells (HSCs) (1), numerous reports demonstrated the ability of the bacteria to invade and remain dormant intracellular to MSCs and HSCs (4–7). We are continuing to explore the role of stem cells in *Mtb* dormancy and reactivation, as well as potential in stem cell niche defense (2, 58, 70). We recently showed that in a mouse model of corona virus infection, dormant *Mtb* reprogram the host CD271+MSC to ASC phenotype which allows the dormant bacteria to undergo reactivation (31). In the present study, in the streptomycin dependent *Mtb*-m18b model of mice infection, we found deep dormant VBNC-*Mtb* infected MSCs in the BAL fluid. We further confirm that these VBNC-*Mtb* infected MSCs reprogram to ASC, and these cells release *Mtb*-Ag (ESAT-6) containing EVs in the BAL fluid. Mice treated with FM19G11, an inhibitor of the ASC phenotype (53) significantly reduced ASCs in the BAL fluid, and associated reduction in the Ag-rich EVs in BAL fluid and the aerosols of these mice. Furthermore, in an in vitro assay, we confirmed the role of VBNC-*Mtb* in MSC to ASC reprogramming, and associated secretion of Ag-rich EVs. These results indicate that ASC mediated secretion of Ag-rich EVs could be part of an innate stem cell defense mechanism against the pathogens’ transmission cycle in to the community. Raghuvansi et al found that MSC suppresses host immune response to help *Mtb* to persist (71). In contrast, our findings suggest that although MSC serves as a host for dormant *Mtb* (1) or MSC reprograms to ASC during d*Mtb* reactivation (31), MSCs also induces an innate defense response characterized by herd immunity i.e. antigen export via EVs for vaccination of contact individuals.

Our study is unique in scientific approach, as we have integrated the metaphysics of a traditional indigenous healing concept, the Nigudah-Ojaskshayamatva or Nigudah-immunity (Figure 1A) to derive our experimental hypothesis on EV mediated antigen-export as a part of immune defense against a pathogen (Figure 6A, and 9C). In this manner, we set an example that the integration of IKS, with modern science may lead to knowledge emergence, an idea that we proposed (14), and working on to discover UPK remain undisclosed/near extinct in rural India. We suggest that the Nigudah-theory of immunity may be the remnant of India’s thousands years of tradition of variolation or inoculation/vaccination against small pox, as mentioned in the Sanskrit text “Sacteya” (32). Interestingly, the skillful application of Kirtan-chanting to spread good-Nigudah into the community may be effective in aerosol inoculation, as recent studies indicate that words including voice plosive consonants emit more aerosol particles from human respiratory tract during speech than words with voiceless fricatives (72). This finding directly correlate with the more aerosol emission during Kirtan chanting as it does such a way, where a constant voice plosive words are played for a longer duration. In this respect, the results of our contact investigation study indicate apparent protection, although it is a small study without enough power to indicate significant protection. However, the efficacy of such antigen-export by aerosols to provide herd immunity remains to be proven. One key challenge is the potential very low amount of antigen that can be delivered through aerosols. Whether this small amount of antigen will be sufficient enough to induce effective immunity need to be investigated.

A previous study showed *Mtb*-antigen export during coughing (69) suggesting that aerosol may spread not only the pathogen but also antigens in the community. Additionally, aerosols may also contain potentially lipoprotein mixtures that may suppress immunity, as *Mtb* was found to use lipoproteins as a decoy mechanism to escape from macrophage response by inhibiting antigen processing and presentation (73, 74). In this context, we found that mouse receiving aerosol-EVs were able to develop immunity, when injected intranasally, and or mouse to mouse aerosol inhalation for two weeks. Hence, it is possible that antigens delivered via EVs may be effective to mount a significant immune response. Infact, Growing bodies of research suggest the important role of EVs in the initiation and maintenance of adaptive immunity (42, 43). EVs are bioactive vesicles formed either as microvesicles (100 nm to >1 uM), exosomes (30-150 nm) or ectosomes (100-500 nm). The role of EVs in adaptive immunity was first noted in three elegant papers published in 1996-1998, where exosomes derived from B cells were found to be enriched for MHC class II, as well as co stimulatory molecules like CD86 and could present antigens to sensitized T cells (75–77), and carriage of tumor antigens to promote antigen-specific T-cell activation (78). Subsequently, gut epithelial cells derived exosomes were found to carry MHC class II/peptides, and involved in dendritic cell mediated immune response (79, 80). Furthermore, exosomes containing viral RNA of infected cells were found to modulate innate immune response (81). In TB, the role of exosomes in cellular immunity has been discovered. *Mtb*-EVs were found to stimulate immunity against *Mtb* suggesting an immune protective activity (82). Then, exosomes of *Mtb* infected macrophage were found to contain antigen capable of activating pathogen specific CD4 T cells in mice (55, 83). These findings indicate that EVs containing *Mtb* antigen mediated immunity may illicit a strong IFN-gamma response. Surprisingly, we failed to isolate EVs secreting macrophages in the BAL fluid of both BCG and AESN-PTB immunized mice. Instead, we identified a CD45-cell phenotype devoid of epithelial cell marker EpCAM but expressing genes involved in MSC derived ASCs. Our results indicate that ASCs exerts an unconventional innate immune response by exporting antigen-packed EVs to aerosols, and thereby vaccinating neighboring animal against *Mtb* infection (61). Khader SA et al showed that vaccination triggered an accelerated interferon-gamma response by Th-17 cells in the lung during subsequent *Mtb* infection (64). Recently, NK cells has been found to be involved in the unconventional immune response of BCG immunized mice challenged with *Mtb* infection (84). Thus, further research is necessary to decipher EV mediated immune response against *Mtb* as an important component of unconventional immune response. One possibility is that EVs illicit a mucosal immunity in the airway mucosa leading to the proliferation of primed T cells, which then migrate to distal lymph nodes and also lung to initiate an innate defense mechanism mediated by ASC. This mechanism may underlie many emerging features of unconventional immune response including NK cells, Th17 cells, as well as hematopoietic progenitors in trained immunity (85–88) (Figure 9C).

We are not yet clear about the source of EVs in the aerosol of smear negative subjects. In the COPD, EVs were found to be produced by airway epithelial cells (89–91) but do not contain MHC-I or II molecules (89, 90). We were not able to confirm MHC expression of isolated EVs because of the limited amount of study material available. The sources of EVs in smear negative subjects may be VBNC-*Mtb* infected macrophages, MSCs or even airway epithelial cells. Or *Mtb* may also be the source, although it is most unlikely, as the isolated EV extract showed the expression of CD63, which is present only in EV of mammalian cells. In an ongoing study, we are looking for VBNC-*Mtb* containing MSC or ASCs in the sputum, as well as BAL fluid of SN-PTB subjects. Findings of the study may provide valuable information about the source host cells of Ag rich EVs in the SN-PTB subjects. Previous studies showed that in the broncho-alveolar lavage (BAL) fluid of treated SN-PTB, a large population of CD4+ T cells is found in flow cytometry that showed reactivity to ESAT-6 (92). In human, about 10% of cells in BAL may be of MSC origin as per a single study (93, 94) Thus, we hope to identify ASCs as well as ESAT-6 reactive CD4 T cells in the BAL fluid of smear negative subjects, and find out any potential association of the presence of these cells with the amount of ESAT-6+ EVs in the aerosols of same subjects. For such an association study, it is also important to use appropriate equipment for the collection and processing of aerosols for the isolation of EVs. We obtained human aerosols after deep expiratory breathing that generates aerosols in bronchioles (95). However, we used either micro spirometer (human) to obtain EBA or a homemade method to collect exhaled breath condensate (EBC), which may not be good enough to collect all the aerosols. In this context, previous studies found that 82% of smear-negative PTB subjects contain ESAT-6-antigen in breath (54), whereas, we found *Mtb* antigen rich EVs only in 70% cases, mainly in the subjects having VBNC-*Mtb* in their sputum. This discrepancy may be related to the fact that we isolated EVs, whereas the study (91) used aerosol pellet directly for ELISA to detect ESAT-6, or that our aerosol collection method is not adequate enough to capture EBA or EBC products. Considering these limitations in human studies, it is important to develop animal model to study antigen-export based mechanism of natural vaccination or herd immunity. In this regard, mouse model described in this study may be of great use.

In summary, we identified an IKS based JivaUpakariTantra approach of aerosol-based inoculation against *Mtb* infection. Underlying principle of the inoculation is the theory of good-nigudah, an ancient concept of microscopic life envisioned by India’s Jain thinkers. A long-term relationship with the IKS through interaction and dialogue led to the emergence of a hypothesis that good-nigudahs are lipid membrane wrapped EVs filled with *Mtb* antigen. Our efforts and results further validate the importance of dialogue and interaction with IKS to enhance the process of knowledge emergence (14). We identified ESAT-6 packed EVs as the good-nigudah being exported in the aerosols of SN-PTB subjects. Our data provide us a strong rationale to develop mouse model of stem cell host/pathogen interaction model to study the possibility that stem cells may have evolved mechanisms to deliver exogenous antigen of pathogens to confer herd immunity against *Mtb*. Additionally, this work highlights the importance of integrated medical education in India for the dialectic interaction between modern and traditional medicine knowledge systems for the advancement of medical science (109).

## METHODS

### Ethical approval

All clinical studies conducted in various institutions were approved by Institutional Ethics Committee of KaviKrishna Laboratory (Contact TB investigation study, and Aerosol collection study of SN-PTB subjects), Guwahati Medical College and Hospital (Aerosol collection study from SN-PTB subjects; protocol: PI: DJK), and RIWATCH, Arunachal Pradesh (Aerosol collection study from SN-PTB subjects), and part of this study data (BM collection) has been previously published (1–2). The animal part of the study was approved by the Institutional animal care and use committee (IACUC) at KaviKrishna Laboratory, Gauhati University, and Forsyth Institute (BCG part of the mice immunization). The biosafety approval (for the in vitro use of BCG and *Mtb*-m18b strain and human BM derived MSCs) was received from the Institutional Biosafety committee at KaviKrishna Laboratory, Forsyth Institute (for the use of BCG, and human BM derived MSCs) and University of Massachusetts-Lowell (for the use of BCG, and human BM derived MSCs).

### Contact TB investigation study

A community based, cross sectional contact TB investigation study was conducted in the village Sualkuchi of Assam with a population of 14000 and very high population density of 3000/km2. Sualkuchi has25 Kirtan temples of close door set-up called, Namghar. In the Hatisatra Namghar, established in 1650 AD, 40-70 devotees between “40-80” ages, both male and female gather daily for two times for community-chanting of Kirtan. Devotee attending this Namghar received regular medical treatment from BD’s clinic as well as JG’s traditional tantric healing during 1993-1994.We selected devotees of HatisatraNamghar to investigate their close contact with a smear positive PTB subject under BD’s treatment; the study was approved and funded in 1994 by Krishna Samaz, the predecessor of KaviKrishna Laboratory (http://www.uniindia.com/~/century-old-family-trust-of-assam-in-revolutionary-cancer-tb-research/States/news/1654921.html). All study subjects were screened and identified as smear positive PTB or smear negative PTB or their close contacts by observing the clinical symptoms for TB, as well as sputum-examination reports obtained from local hospital (96). The study was implemented by field based contact investigation including home visit, Namghar visit and then confirmed with their concerned physicians and health workers. The 8 close contacts of the index smear positive case that didn’t attend Kirtan chanting was of <10 years and thus suggested for further clinical evaluation. This cross-sectional study was repeated in 1998, and again in 2009. From 2010-2019, study was converted into a longitudinal study, and the details of the study subjects were maintained in lab register using a unique identity number, followed up with regular interval by the Physician assistants and monitored by the Supervisor. The Supervisor visited the study subjects at least twice in a year during the period of the contact investigation study. The study was supervised and conducted by BD with his physician assistant team (LP, TB, PS, ND and RD) and the activities performed were standard TB care protocol. Thus, no informed consent was required from the participants. Local administrative approvals were obtained from the chief of the Namghar and later study was also ethically approved by Institutional Ethics committee, KaviKrishna Laboratory.

### Suppositional reasoning (tarka) based modeling of good-nigudah

Suppositional reasoning (“tarka”) is a well-established investigation method for new knowledge generation in ancient Indian philosophical system (97) (details in supplementary note 1). Tarka emphasizes on the social dimension to knowledge, where reasoning reigns resolving controversy in ways over and above the sources and facilitate the process of knowledge-emergence process within an IKS (14). The IKS of JivaUpakarTantra or Vedic Altruism has evolved an unique tarka system, the satvata (examination of truth) tarka; Krishna Ram Das (KRD) used this tarka to enhance his Anviksiki (logic based investigation) skills for composing Kamrupi style of poetry (98), whereas JG used the tarka to enhance tantric healing practice (Chapter 2.2, pages 42-47, Reference 17). The tarka has five major components: observation of a desire (ichha-prataskhya), inference based on prior knowledge (satvata/purvavat-anuman), suppositional reasoning (yukti-tarka), special feature of doubt (samasya-laksana) and examination of a new idea (parkisa-dharana); some of these five elements were part of an ancient system of knowledge generation through reasoning (https://plato.stanford.edu/archives/sum2019/entries/early-modern-india/ (37), whereas the other elements are the product of local transformation of knowledge system as predicted by local adaptation of indigenous knowledge system (15, 30, 99). JG and KRD used satvatatarka tool to convince BD to do the contact-investigation study among the temple devotee, whereas Dayal Krishna Bora (DKB), an eminent Vedic scholar convinced BD to use the tarka method as a tool for the metaphysical modeling of good-nigudah. DKB was familiar with JG and KRD’s Satvatatarka method, as the trios were involved in researching on JivaUpakarTantra (page 53, 428-530, Reference 98). Thus, DKB represented the good-nigudah-hypothesis side, whereas BD represented the modern medicine side (two circles of tarka, Figure 4A, and supplementary note 1). BD started the debate with an initial enumeration (udeshya): the devotee who was chanting Kirtan did not suffer from PTB, because they were infected by the bacteria; there body developed immunity against the pathogen. DKB identified the element of anista-prasangayukti (reasoning based on unwanted consequence) in BD’s presumption. DKB stated that our conclusion that devotee may be infected by the pathogen is an unwanted effect of BD’s initial observation that smear-negative subjects do not show bacteria in sputum; without bacteria in their sputum, how could they spread the pathogen to others? BD replied that 30% of smear-negative subjects convert to smear-positive subjects as the disease progress. So, it is possible that 3 of the smear-negative chanting devotees eventually became smear positive and infected the rest, who received low-dose of bacteria enough to trigger immunity to resist bacterial growth in lung. Those who did not attend Kirtan did not get infected with a low dose bacterium, and therefore, when they were exposed to high dose bacteria by smear-positive people in the community, their immunity was not strong enough to resist the bacterial growth in the lung. However, DKB reached an alternative explanation by applying the purvavat-anuman method, which is the unperceived effect from a perceived cause (37)(page-32). DKB inferred that good-nigudahs spreading from the smear-negative subjects during Kirtan chanting protected the rest of the devotees from TB. To this explanation, BD applied samasya-laksana (doubtful feature) and stated that good-nigudah is an imaginary thing, and therefore, cannot be a factor for reasoning. To this argument, DKB applied satavata-anuman (inference based on prior knowledge), and referred to an ancient law of duality (dvaita-vedanta), where it is possible for good to emerge from bad as per Avatar doctrine (Chapter 1.4, page 14; Reference 17). DKB further elaborated how good can arise from evil by applying another sutra (theory), the protibimbawada (theory of reflection) by a renowned scholar Padmapada (110). DKB asked BD to draw the general features (laksana) of the TB causing bacteria in a piece of paper. BD drew a seed (DNA and cytoplasm) inside a sack (plasma membrane), Figure 4B. DKB then applied the theory of reflection to draw a similar sack with part of the seed inside, and termed it as good-nigudah. DKB then explained that as per the law of duality (dvaitavada; https://en.wikipedia.org/wiki/Dvaita_Vedanta), some good chemical exist in the bacteria, which will be present in the seed of good-nigudah. DKB then asked BD to come up with any known features by medical science that can resemble the image of good-nigudah (Figure 4B). To this question, BD replied that bacteria-infected cells of our body may be able to capture some antigens from the bacteria, and put them inside a vesicle and secrete them outside; such vesicles are known as extracellular vesicles (EVs). DKB then invoked the observation of a desire (ichha-prataskhya) element to visualize a form of good-nigudah, and BD drew it near the image of bad-nigudah (Figure 4B). Both DKR and BD then invoked the last element of the tarka, pariksha-dharana or the examination of a new idea. BD then hypothesized that an extracellular vesicle-containing antigen derived from *Mtb* may serve as the model for good-nigudah. The satvata-tarka session ended.

### The clinical study of aerosol and sputum collection from smear-negative PTB subjects

The clinical study was approved by “Institutional Ethics Committee” of KaviKrishna Laboratory and KaviKrishna Telemedicine Care, a branch of KaviKrishna laboratory, and also RIWATCH. Smear negative subjects were selected as described previously (1, 67) and in the result section. The mean age of the study group was 46 years (range 42 to 75 years), and the ratio of male to female participants were 12:14. Some of the subjects had hypertension (n=8), and suspected with chronic-obstructive pulmonary disease (COPD) (n=12) under occasional treatment with antibiotics and bronchodilators for last 4-5 years. None of the study subjects had prior history of diabetes, cancer and any history of extra-pulmonary tuberculosis. These patients were smear negative PTB cases (no *Mtb*-DNA in sputum; bacterial culture was negative) and were being prescribed with a course of anti-TB drugs by the district TB control program. Briefly, aerosol samples were collected from 18 smear negative PTB subjects under anti TB treatment (8/18) or not yet started the treatment (10/18) by exhaled aerosol breath (EBA) method. These subjects were prepared (asked to take rest for 2-3 hours before the procedure, drink enough liquid, and then do not take any food/drink an hour before the procedure), and asked to exhale into a micro-spirometer after a deep breath so that their forced expiratory volume can reach the FEV1 (forced expiratory volume in one second)>2 liters and PEF (peak expiratory flow) of > 300 liters/minute. They were then asked to exhale by mouth into the spirometer mouthpiece for every 3-5 seconds. The breathing process was done for 10 minutes. The procedure was well tolerated by all the participants. The spirometer was taken into the laboratory for the aerosol collection, and processing for the isolation of extracellular vesicles (EVs) (details are explained in the method section for extracellular vesicle isolation). The aerosol samples (suspended in 0.1% BSA) were examined for amylase activity (#ab102523, Abcam; detection limit of 0.2 umol/l) and found to contain <1 umol/l amylase suggesting minimal contamination by saliva. The protein content in the aerosol was measured by Bedford assay (micro assay; detection limit of >1 μg /ml); we found average 10.4 ± 4.2μg/ml protein per 15 minutes of procedure, out of which, 2.5 ± 1.4 μg/ml were EVs. To increase the EV yield and also make the sample saliva free, we have used the exhaled breath condensate (EBC) method of aerosol collection previously used to isolate EVs in subjects with COPD (49). Thus, in selected subjects (where ESAT-6 EVs was detected following initial screening, and required enough EVs for in vivo mouse experiments), a home based EBC collection method was used. Briefly, subjects were prepared 3-4 hours before the procedure, and asked to exhale into a drinking glass (250 ml size). The drinking glass was soaked in ice cold 0.2% BSA, remaining BSA was discarded, and then the glass was wrapped in a frozen gel pack. Study subjects were asked to exhale into the glass for 15-20 minutes; frozen gel pack was changed periodically 4-5 times to keep the glass ice cold. EBC (approximately 1.5 ml containing 20.5 ± 6.3 μg/ml protein, out of which, on average 5 μg/ml was EVs; no amylase was detected suggesting minimal contamination with saliva) was collected and immediately subjected for EV separation (described below). The procedure was repeated 3-4 times weekly for 2-3 weeks to obtain enough EV materials (50 μg/subject containing >10μg of ESAT-6, enough doses to immunize 5 mice) for the mouse experiments (Figure 6-7). Then, collected EVs were concentrated in a 0.5 ml aliquot in PBS containing 0.1% BSA, and stored at −80C for long-term storage for in the in vivo use as previously described (100). Sputum samples were also collected from these subjects over three consecutive days, and then transported to Ranbaxy lab, Mumbai following RNTCP guidelines for culture and sensitivity test. Simultaneously part of the sputum samples were subjected to a liquid culture method that facilitates resuscitation of difficult to culture or VBNC *Mtb* at KaviKrishna lab (1) using early stationary phase supernatant (ESPSN) of *Mtb*-H37Ra, a conditioned media that can resusciate viable but non culturable *Mtb* (65, 101). The *Mtb*-CFU report for the sputum samples from Ranbaxy lab was negative. The +ve liquid culture (as confirmed by AFB staining) following ESPSN treatment was identified to contain VBNC-*Mtb*.

### Isolation of extracellular vesicles (EVs) from aerosols from patient subject and mice

We have had extensive experience of isolating cell membrane fraction using ultracentrifugation (47), which we utilized for the isolation of EVs from aerosols. In order to collect the putative *Mtb*-antigen rich EVs in the exhaled breath aerosol (EBA) of the smear-negative individuals, the spirometer mouth piece and tube were rinsed with 5 ml of PBS containing 0.1% BSA. A cell-free supernatant was obtained by passing the 5mL BSA solution containing aerosol through a 0.45-mm–pore size polyvinylidenedifluoride filter (Millipore). A similar technique was used to collect the exhaled breath condensate (EBC). The supernatant thus obtained was concentrated approximately 20-fold using an Amicon Ultrafiltration System (Millipore) with a 100-kDa exclusion filter. The concentrate was then stored at −80°C until further use. Then the concentrated supernatant was ultracentrifuged at 110,000 g for 1 hour at 4°C (Beckman Coulter L100k, company name) to sediment the vesicular fraction into a pellet. The supernatant was then discarded; the pellet was collected in 0.1% BSA to make the EV solution and stored at −80°C. This is how the ultracentrifugation method of EV collection was accomplished using a modified protocol that we developed from our experience in cell membrane isolation (47), which is similar to established techniques (102). For the ELISA study, EVs were suspended in RIPA buffer (100 µl EVs was mixed with 11 µl of 10x RIPA (47), and incubated at 5C for 5 minutes) and a cocktail of protease inhibitors (1 mM phenyl-methylsulfonyl fluoride and 1 ug/ml leupeptin) to obtain a protein lysate (12) and then subjected to ELISA study for ESAT-6 antigen or CD63, a marker of EVs. In this way, an initial screening of 18 subjects was achieved to select the subjects having highest level of ESAT-6+ EVs. To rule out false positive ESAT-6 EVs, EVs of aerosols were also obtained by the direct immunoaffinity or immunomagnetic sorting of CD63+ EVs as described (102). Here, we used EBC derived aerosols from 3 SN-PTB subjects from 18 initially screened subjects for ESAT-6 EVs (patients showing highest, lowest and moderate ESAT-6) to obtain better yield of EVs (49). For this purpose, we first prepared the CD63-magnetic beads by using a protocol that we extensively used to prepare CD271-magnetic beads (2, 12). For this purpose, a CD63 antibody (#NBP2-42225; Novus Biologicals, CO) was first PE conjugated by SiteClick antibody labeling kit (#S10467; Life Technologies, Grand Island, NY), and then an Easy Sep PE sorting kit (#18554; Stem Cell Technologies, Vancouver, BC) was used to perform the positive selection of CD63+ EVs in the ultrafiltration (110 kDa exclusion filter, Millipore) product. This CD63+ sorted EVs were subjected to western blotting to confirm the presence of ESAT-6 antigen. As EBC derived aerosols showed high and better yield of EVs than EBA, EBC was performed for all the study subjects and EVs were isolated from EBC derived aerosols by ultracentrifugation and stored at −80 for future animal studies.

The EVs were also isolated from the aerosols of immunized *Mtb*-m18b infected mice (n=5 for each of the four subjects). Briefly, *Mtb*-m18b infected immunized mice (8 weeks after infection) were allowed to breath inside a 50ml tube for 5 minutes/day following sneezing-induction by intranasal injection of hypertonic saline (10 μl drop once that cause 4-5 sneeze/ 5 minutes). PBS was added in the tube where aerosols were collected from 5 mice (total 50 minutes of sneezing in two weeks). Aerosols were processed for ultrafiltration followed by ultracentrifugation to obtain EVs as described above. Approximately ∼15 ug EVs were collected/5 mice/50 minutes of cumulative sneezing in two weeks.

### Dynamic light scattering

The size distribution of EVs was measured on a ZetasizerNano (Malvern, United Kingdom) at 20°C using an angle of 173 degrees and 633-nm laser.

### Transmission Electron Microscopy

Visualization of EVs was accomplished by resin-embedding method of transmission EM. Selected samples of ultracentrifugation product were fixed 4% paraformaldehyde, postfixed in 1% osmium tetroxide (OsO_4_) for 25 min followed by rinsing twice in distilled water. Then, the fixed EVs were dehydrated by ethanol and subjected to block staining with 1% uranyl-acetate in 50% ethanol for 30 min and embedded in Taab 812 (Taab). Then, ultrathin sections were obtained and analyzed with a JEOL JEM1400 (JEOL USA Inc, Peabody, MA) electron microscope equipped with Gatan orius, a 10.7 megapixel side-mounted transmission EM CCD camera.

### Enzyme Linked Immunosorbent Assay (ELISA)

Sandwich ELISA was used to check the presence of ESAT-6 antigen on the EVs isolated from the aerosols of smear negative PTB patients, aerosols, broncho-alveolar lavage (BAL) fluids, CD45-/CD45+ cells of BAL fluid of m18b infected immunized mice as well as FM19G11 treated m18b infected immunized mice and conditioned media of ASCs (m18b derived VBNC infected ASC with/without FM19G11). Samples were dissolved in RIPA buffer and processed for In-cell ELISA. Briefly, 96-well ELISA plates were coated overnight with 1 µg/well of polyclonal rabbit anti-ESAT-6 Ab (#ab45073, Abcam Inc, USA) followed by blocking with 2% BSA for 1 h at 37° C or overnight at 4° C. EVs suspended in RIPA was centrifuged, and 50 μl of supernatant was added to the wells of the ELISA plate, covered and incubated overnight at 4°C. Captured ESAT-6 antigens were detected with mouse anti-ESAT-6 Ab, clone 11G4 (#ab26246, Abcam Inc, USA), diluted 1∶150 and HRP conjugated anti-rabbit Ab (Sigma-Aldrich, USA), diluted 1∶1000. A similar sandwich ELISA was used for detection of Ag85, where the ELISA plate was coated with 1 µg/well rabbit-anti-Ag85 Ab (#ab43019, Abcam Inc, USA) as captured antibody, and mouse anti-Ag85 Ab (#ab36731, Abcam Inc, USA), diluted 1∶1000 was used as detection antibody. All Ab dilutions were made in blocking buffer (1× PBS+2% BSA). At each step, ELISA plates were thoroughly washed 2 times each using 1× PBS+0.05% Tween 20 (AMRESCO, USA) and 1× PBS. The bound HRP was detected using 100 ul of 3,3’,5,5’-tetramethylbenzidine (TMB) substrate for each well followed by 20 minutes of incubation at room temperature; the reaction was stopped by the addition of 0.1 N Sulfuric acid. The absorbance was measured at 450 nm. For the detection of ESAT-6 in mouse BAL fluid derived EVs, the rabbit anti-ESAT-6 (#ab45073, AbcamInc, USA) was used as capture antibody. The CD63 ELISA was performed using the same protocol; a mouse monoclonal CD63 antibody (#sc-5275, SantaCruz Biotechnology, Dallas, Texas, Novus Biologicals, Centennial, CO, USA). For the quantification of ESAT-6 protein in EVs, a standard curve of absorbance value was obtained using 1-500 pg/ml concentration of recombinant ESAT-6 (NBP1-99045; Novus Biologicals, Centennial, CO).

### Western blot

The procedure was performed as described previously (12). The EVs pellet suspended in RIPA was used to perform western blot to check the expression of ESAT-6 (anti-ESAT-6 antibody (#ab26246, AbcamInc, USA) and CD63 (#NBP2-42225SS, Novus Biologicals, Centennial, CO, USA).

### Aerosol-EV induced ESAT-6 specific IFN-gamma response in mice

All the necessary experimental procedures were undertaken in accordance with approvals of Institutional Animal Ethics Committee and Institutional Ethics Committee of KaviKrishna Laboratory. The 6-8-week-old C57BL/6 female mice were obtained from National Institution of Nutrition, Hyderabad, India and were maintained in the animal house at pathogen free condition as previously described (1). We have used EVs obtained by ultracentrifugation from EBC derived aerosols of all the study subjects after confirming the presence of ESAT-6 and CD63 by ELISA. Then the mice (C57BL/6, 6-8 weeks, female) were twice (a week apart) challenged with intranasal injection of 0.5 μg/ml (20 ul in 0.1 BSA) aerosol-EVs of all study subjects. Dormant infection was achieved by *Mtb*-m18b infection in mice (10^6 CFUs i.v. and 2 mg streptomycin/daily/two weeks) followed by two-months of streptomycin withdrawal (59). After 6 weeks, splenic cells were obtained and challenged with either ESAT-6 (10 μg/ml) or Ag85 (10 μg/ml), and then subjected to IFN-gamma measurement by ELISA. The smear negative PTB (SN-PTB) subjects 2-5 immunized mice were selected for m18b *Mtb* infection and herd immunity mice model, as their IFN response against ESAT-6 was >1.5ng/ml, a value that can resist bacterial invasion (59).

### Mice Model for Herd Immunity

We tested whether immunization with aerosol EVs served as a protective mechanism in mice against a *M. tuberculosis* challenge as described earlier (82); we used the *Mtb*-m18b strain (1) and the mice immunized with BCG, aerosol-EVs of SN-PTB and COPD subjects. We have selected 2-5 AESN-PTB subjects immunized mice for m18b infection, as their IFN response against ESAT-6 was >1.5ng/ml, a value that can resist bacterial invasion (59). Briefly, mice immunized with BCG or aerosol-EVs of smear-negative PTB subjects and COPD were infected with approximately 100 m18b-CFU via aerosol, and then treated with streptomycin for three weeks to establish lung infection (1, 59). Six weeks after infection, lung bacterial burdens were determined by *Mtb*-CFU assay. Mice immunized with PBS or BCG Pasteur strain (5×10^5 CFU in 30 μl PBS/mouse/intra nasal once) were used as negative and positive controls, respectively. The experiment was done four times with aerosol of four different subjects. Also, we investigated, if the aerosol-mediated immunity against *Mtb*-m18b infection also involves antigen-export to neighbors, we developed a unique mouse model where aerosols from infected mice were allowed to be inhaled by healthy mice. The healthy mice were exposed to the aerosol of m18b-infected mice that received prior immunization either with BCG or Aerosol-EV of SN-PTB subjects. Briefly, the m18b infected mice (4 weeks after infection) were allowed to breath inside a 50ml tube for 5 minutes/day following sneezing-induction by intranasal injection of hypertonic saline (10 μl drop once that cause 4-5 sneeze/ 5 minutes); immediately after, healthy mice were allowed to breath into the aerosol rich tube for 5 minutes/day. The procedure was repeated for two weeks, and then both infected mice and healthy mice were sacrificed. Then, the healthy infected mice were subjected to measure *Mtb*-CFUs in their BAL fluid, and whole lung to confirm the immunization against m18b infection.

### Isolation of BAL fluid from mice

The BAL fluid from the *Mtb*-infected mice was collected as previously described (103). Briefly, mice were euthanized by cervical dislocation and a catheter of about 0.5cm was inserted into the trachea and a syringe loaded with a balanced salt solution with 100 µM EDTA was connected to the catheter. The salt/EDTA solution was gently injected into the lung and is aspirated back gently. The above step was repeated more than twice and obtained 2 ml of BAL fluid (103). This BAL fluid was further used for isolation of CD45+/CD45-cells followed by detection of ESAT-6 and *Mtb*-CFU.

### *Mtb*-CFU assay

The *Mtb*-CFU in lung, BAL fluid and CD45-/+ cells of BAL fluid obtained from immunized m18b infected mice were counted as previously described (1, 2). The intracellular and extracellular *Mtb*-CFU of CD271+BM-MSCs infected with m18b derived VBNC-*Mtb* were detected as previously described (31). The immunized mice after 8 weeks of m18b infection followed by BAL fluid and aerosol isolation were sacrificed. The lungs and spleens of the sacrificed mice were removed aseptically and were homogenized in PBS with 0.05% Tween 80, and then subjected to CFU assay (2). For mouse tissues infected with m18b, *Mtb*-CFU media was supplemented with 50µg/mg streptomycin as described (1).

### Identification of viable but non culturable *Mtb* (VBNC-*Mtb*)

The CD45-cell intracellular *Mtb*-m18b-GFP bacteria were manually counted per microscopic field, and the data was used to find number of bacteria per 1×10^4 cells. The data was compared with the *Mtb*-DNA copy number, and *Mtb*-CFUs with and without early stationary phase supernatant (ESPSN) of *Mtb*-H37Ra, a conditioned media that can resuscitate VBNC-*Mtb* (65) and thus used to identify the VBNC in samples. Thus, VBNC was identified when we find that *Mtb* DNA copy number value is higher than *Mtb*-CFU value and *Mtb*-CFU value with ESPSN is higher than without ESPSN.

In vitro VBNC production from m18b strain, *Mtb* are first exposed to chronic hypoxia to obtain non-replicating persister II (NRP II) dormant *Mtb*; these dormant *Mtb* were co-cultured with macrophages for 48 hours under extreme hypoxia followed by reoxygenation. As a result most of the NRP II converts to VBNC. CD271+ BM-MSCs are capable of resuscitating these VBNC in an in vitro co-culture assay for two weeks.

To confirm the presence of VBNC in the sputum samples of smear-negative subjects (n=10), sputum was decontaminated using 4% sodium hydroxide and kept in the BACTEC tube that contain broad spectrum antibiotic cocktails for 72 hours, which mostly kill the contaminated bacteria living behind *Mtb*. This processed sputum was then diluted in PBS, centrifuged and then added to MSC cultures for 8 hours and then cells were washed and treated with amikacin and metronidazole to avoid contamination; we used a protocol that we developed for culturing cancer cells with saliva/sputum (104). As a positive control for VBNC detection, processed sputum were allowed to grown in 7H9 liquid supplemented with prepared 50% (vol/vol) resuscitation factor EPSN along with 10% (vol/vol) oadc (oleic acid, albumin, dextrose, and catalase; bd biosciences), 0.05% (wt/vol) tween 80 or on 7H10 agar plates. EPSN comprises of sterile culture supernatant of *M. tuberculosis* H37Ra standing culture (the optical density at 600 nm was about 1.2) grown in 7H9 medium. Culture supernatant was collected, centrifuged at 6,000 3 g following sterilization by filtration through a 0.22-mm-pore size filter (101). After two weeks, both the MSC and EPSN cultures were evaluated for *Mtb*-CFUs. For the MSC culture, the conditioned medium of the MSCs was collected and further used for EVs isolation and detection of ESAT-6 detection in the EVs as described above. The MSCs were lysed and subjected to colony-forming unit (CFU) assay as described previously (1). Briefly, pellet was lysed with 1 ml of 0.1% Triton X-100 for 15 min, vortexed for 30 seconds, and serial 10-fold dilution was prepared in Middlebrook 7H9 broth. The diluted cell lysate was then placed separately on 7H10 agar plates (BD Bioscience, #295964). All dilutions were made in quadruplicate. The cultures plates were incubated at 37°c for up to 8 weeks and CFUs were counted for all subjects with/without ESPSN. CFUs were plotted as the means of log10 CFUs per mL of the sputum.

### Assay for VBNC induced MSC to ASC reprograming

The ASC reprograming of VBNC infected MSCs was confirmed as previously described (31, 105). Briefly, human CD271+ BM-MSCs were obtained from healthy volunteers after informed consent (1) and infected with VBNC of *Mtb*-m18b. The infected BM-MSCs were grown in serum free media for naïve MSCs (58) for two weeks; during this time, cells were recovered and subjected to In Cell ELISA to evaluate p53/MDM2 oscillation, and the protein level of active caspase-3. After two weeks of in vitro culture were collected and subjected to real time PCR gene expression study to confirm ASC reprograming To isolate the mRNA from VBNC infected CD271+MSCs, we used the µMACSTM technology (Miltenyi Biotec, Bergisch-Gladbach, Germany) as described previously (1). Briefly, the mRNA is isolated in a magnetic column using super paramagnetic Oligo (dT) Microbeads, which target the poly RNA tail of mammalian RNA. The mRNA gets attached to the magnetic column, whereas the bacterial and stem cell DNA remained in the passed-through lysate. As per manufacture instruction, the mRNA was converted to cDNA in a heated magnetic bar and was subjected to qPCR analysis using the TaqMan Gene expression assay Biotech, Bergisch-Gladbach, Germany). RNA quantification was done using the ΔΔ Ct method by using the SDS software, version 2.2.1 (1). The following TaqMan primers were used: Human: CD271 (Hs00609976_m1), CD45 (Hs00898488_m1), CD44 (Hs01075861_m1), CD133 (Hs01009250_m1), CD34 (Hs00990732_m1), HIF-2alpha (Hs01026149_m1), HIF-1alpha (Hs00153153), ABCG2 (Hs01053790_m1), Oct-4 (Hs03005111_g1), Nanog (Hs02387400_g1), SOX-2 (Hs01053049_s1), CD73 (Hs01573922_m1), p53 (Hs00251516_m1) and p21 (Hs00355782_m1).

### Isolation of CD45-cells in BAL fluid

Accordingly, mononuclear cells of BAL fluid were pooled from 10 mice to collect enough cells (average 1×10^6 cells). Isolated cells were subjected to immunomagnetic sorting for CD45+ (enriched in macrophages) and CD45-cells (enriched in MSCs) as described previously (2). Sorted cells were then in vitro cultured for 1 day using 0.4 micron cell insert (106) to collect bacteria free secretory products in the conditioned media (CM). The cells on the top of the insert was lysed, and subjected to *Mtb*-CFU assay, whereas the CM was subjected for EV enrichment, followed by ELISA assay to detect the presence of ESAT-6 antigen.

To evaluate the ASC phenotype of *Mtb* harboring CD45-cells, we performed an experiment where bacterial DNA and host cell RNA can be simultaneously obtained (1). The CD45-cells recovered from BAL-fluid were subjected to immunomagnetic separation of mammalian RNA and bacteria DNA. The RNA was used to evaluate the ASC phenotype of CD45-cells by performing gene expression of stemness genes expressed in ASCs (31). As per manufacture instruction, the mRNA was converted to cDNA in a heated magnetic bar and was subjected to qPCR analysis using the TaqMan Gene expression assay Biotech, Bergisch-Gladbach, Germany). RNA quantification was done using the ΔΔ Ct method by using the SDS software, version 2.2.1 (1). The following TaqMan primers were used: Mouse: CD271 (Mm 00446296_m1), CD45 (Mm 01293575_m1), CD44 (Mm01277163_m1), HIF-2alpha (Mm01236112_m1), ABCG2 (Mm00496364_m1), Oct-4 (Mm00658129_g1), Nanog (Mm02019550_s1), Sox-2 (Mm03053810_s1), HIF-1alpha (Mm00468869_m1), p53 (Mm01731290_g1), p21 (Mm01332263_m1), MDM2 (Mm01233136_m1), Glutamate Cysteine Ligase (GCL) (Mm00802655_m1), CD73 (Mm00501915_m1), CD11b (Mm 00434455_m1) and GAPDH (Mm99999915_g1).

### Immunofluorescence assay

Sorted CD45-cells of BAL fluid (from AESN-PTB subject #2; on day-90 after aerosol-EV immunization, infected with *Mtb*-m18b-GFP) were cultured in the glass chamber coated with polylysine (10ug/ml/overnight at RT), air dry in laminar flow hood, and then add fibronectin (10ug/ml) for overnight at 37c, remove it, wash with 0.1% BSA, and add cells immediately without letting it dry so that cells will stick and within next 24-48 hours have taken images.

### *Mtb*-DNA copy number analysis

Assay was performed as described previously (1–2). Bacterial DNA was isolated using the μMACSTM technology (MiltenyiBiotech, Bergisch-Gladbach, Germany). The exhaled breath aerosol (EBA) of the smear-negative PTB subjects collected in the spirometer was rinsed with PBS and was analyzed for the presence of *Mtb*-DNA. 150 *Mtb*-m18b were pulsed to an EBA sample (10 μg/ml protein in 100 ul PBS), which served as a positive control. Lungs and spleen of mice immunized with *Mtb*-m18b, BCG, AESN-PTB, and aerosol-EVs obtained from COPD were also tested for the presence of *Mtb*-DNA. *Mtb*-DNA was isolated using the manufacturer’s instructions. To amplify the *Mtb*-DNA a *Mtb* specific TaqMan primer (#rep 13E12, Quantification of Mycobacterium Tuberculosis, Advanced Kit; Primer Design Ltd, Southampton, Hants,UK) was used as described earlier (1), and a *Mtb*-DNA standard curve. The qPCR was performed in a 7900 system (Applied Biosystems, Foster City, CA) and *Mtb* copy number was obtained from the standard curve as per instructions provided in the Primer Design Kit.

### In Cell Western Assay for aerosol EVs immune response in DC

This was carried out as described earlier (53). Briefly, bone marrow mononuclear cells were collected from a mouse femur (∼2 × 10^7^ cells) into 2 ml of Iscove’s Modified Dulbecco’s medium supplemented with 2% FBS as previously described (114). 1 × 10^6^ cells were plated in a 48-well plate per well in RPMI with 10% heat-inactivated FBS. This was followed by induction with GM-CSF (Invitrogen) 5 ng/ml for 4 days which helps in the differentiation of BM-MNC to dendritic cells. The floating cells were removed and 2.5ng/ml GM-CSF along with 0.5 ug/ml EVs in 20 ul of 0.1% BSA was added to the adherent cells for 4days. On day-8 the cells were fixed, and In-Cell Western was carried out as per manufacturer instructions (Licor Bioscience, Lincoln, NE, www.licor.com). After fixation, the plates were washed with PBS and MHC II and CD11c primary antibodies were added in 2% BSA and 0.1% saponin. After overnight incubation of primary antibody incubations at 4°C, plates were washed thrice (5 minutes each) with PBS-T (PBS with Tween 20) at RT. Then, secondary antibodies conjugated to IRDye 800CW (goat-anti-rabbit-IgG (Licor Biosciences, NE) or goat-anti-mouse-IgG (Licor Biosciences, NE) was used. Plates were then incubated with secondary antibody solutions for 90 minutes at RT in the dark. After secondary antibody incubations, plates were washed thrice times (10 minutes each) with PBS-T at RT in the dark, and air-dried before scanning. Plates were scanned and analyzed using an Odyssey IR scanner using Odyssey imaging software 3.0.

### Statistics

Statistical analysis was carried out using GraphPad Prism version 4.0 (Hearne Scientific Software, Chicago, IL, USA). Student’s t test was used for comparisons with Newman-Keul post hoc test for the *Mtb*-CFU assay, and ANOVA withDunnett post hoc test was performed for the rest of the experiments. Data are expressed as means ± SEM; *P < 0.05, **P < 0.001, ***P < 0.0001

## Abbreviations

SN-PTB: smear-negative Pulmonary tuberculosis
AESN-PTB: Aerosol derived extracellular vesicles of smear negative PTB subject
SP-PTB: smear positive pulmonary tuberculosis
FGD: focused group discussion
JUT: Jiva Upakar Tantra (Vedic Altruism)

## Acknowledgement

We thank Idu-Mishimi people of Arunachal Pradesh, India, and the Vaishnav devotee of Hatisatra (Sualkuchi, Assam, India) including late Soneswar Atoi for support and co-operation while conducting the clinical part of the study. Special thanks go to Prof Gyi of Mongar Referral Hospital of Bhutan for fruitful discussion during the early phase of the contact investigation study during 1994-1998. We thank staff of animal lab facilities at Forsyth Institute, Gauhati University and KaviKrishna laboratory for their support in conducting the BCG, and *Mtb*-m18b related in vivo work. Funding: We thank Dieter Beer and the family of Toronto, Canada for continuing financial and logistic support since 1998 to present for the completion of this study.

## Additional funding

KaviKrishna Foundation, Assam, India (BD, LP,SG, PS, TB, NRD, RD, MM, SB, JT, SS, DK) Laurel Foundation, California, USA) (BD) and KaviKrishna USA Foundation, MA, USA (BD, BP and PG), and Department of Biotechnology, Government of India (BT/PR22952/NER/95/572/2017) (BD, IP, LP, PS, TB).

## Author’s contribution

B.D., JG and KRD initiated the study. B.D. designed and supervised the study. B.D. D.K, I.P., L.P., T.B., N.D., M.M., P.S., R.D., J.T., P.G., and R.M. performed contact investigation as well as clinical part of the study. B.D., J.G., K.R.D. and D.K.B. conducted the metaphysical modeling part of the study (Figure 1 and 4). B.D., L.P., B.P., S.B., S.S. and P.S. performed laboratory part of the study. B.D., L.P., B.P., S.G., D.K. and I.P. analyzed the data. B.D., L.P., and S.G. wrote the paper.

## Supplementary Note 1: Jiva Upakar Tantra (JUT) or Vedic altruism based Satvata tarka, and Prajna Agan (Knowledge emergence)

### Background: Jiva Upakar Tantra or Vedic Altruism

Jiva Upakara Tantra (JUT) is an a traditional medical text of ancient Kamarupa (Sualkuchi-Hajo cultural complex located 25 miles north east of Guwahati, the main city of Assam, India). Although the text was lost, a few practitioners maintained the healing tradition, and a poet-philosopher Krishna Ram Das (KRD) revived the philosophical aspect of JUT by writing several poems (Sonali Nakhar Jui or Flame of Golden Nail; publisher Assam Book Depo, Guwahati, 1991. Reprint 2019). The key philosophy of JUT is the “Avatar Kosha” (Avatar layer), a transient subtle body that emerges in our body during sickness, similar to the emergence of Avatara such as Fish Avatar during the Great Flood. Because, Avatar acts altruistically (Upakara in Sanskrit), hence, KRD translated Jiva Upakar Tantra as Vedic Altruism, the Vedic doctrine of altruism (Chapter 2, Reference 17).

Bikul Das (BD), the lead author of this paper did research on JUT both in Assam and also adjacent Bhutan between 1995-1998. BD found JUT to be a form of Indigenous Knowledge System, and borrowed its philosophical idea, Prajna Agan (equivalent to western idea of Knowledge emergence) to develop a concept of cultural emergence (Das B. Globalization and Emerging Opportunities of Indigenous Cultures. Proceeding of the first annual conference of International Society for Cultural Studies, Mumbai, India 2003. https://www.academia.edu/7882695/Globalization_and_Emerging_Opportunities_for_Indigenous_Cultures).

IKS is the knowledge system inherited by indigenous culture, and it operates at local level of interaction through tradition/culture, and exhibit emergent property i.e. capable of producing new knowledge through interaction, and adaption to local environments (14, 15).

As depicted in Figure 1, JUT has a unique metaphysics of medical treatment. The metaphysics is based on an Ayurvedic concept of “Vyadhikshamatwa” (equivalent to immune defense mechanism) (34–36) that body defense mechanism of a TB subject can extract good-nigudah (mentioned in ancient Hindu and Jain sacred texts) from bad-nigudah (pathogen itself as per JUT-UPK) of PTB subjects and spread these extracted good-nigudha via aerosol inoculation during Kirtan chanting to devotees. Then, this good-nigudah provides protection to the community against bad-nigudah spread via aerosol from other PTB subjects. In JUT, the body defense mechanism of good-nigudah is known as the “Nigudah-Ojaskshayamatva pravriti-arthapati”, which can be roughly translated as the hypothesis (pravriti-arthapati) of “Nigudah-immunity”. The immunity activates a subtle layer, the Avatar Kosha (Altruistic layer of subtle body), and this is why the treatment method is known as Jiva Uapakar (Biological altruism). Since the healing method has its root in the Vedic concept of Avatar (around the cultural complex of Hayagriva Temple of Hajo, Kamarupa), hence, it is also known as Vedic Altruism (Chapter 1-3; Reference 17). This idea of Avatar Kosha is based on the Satvata tarka element as described below.

**Figure 1:**
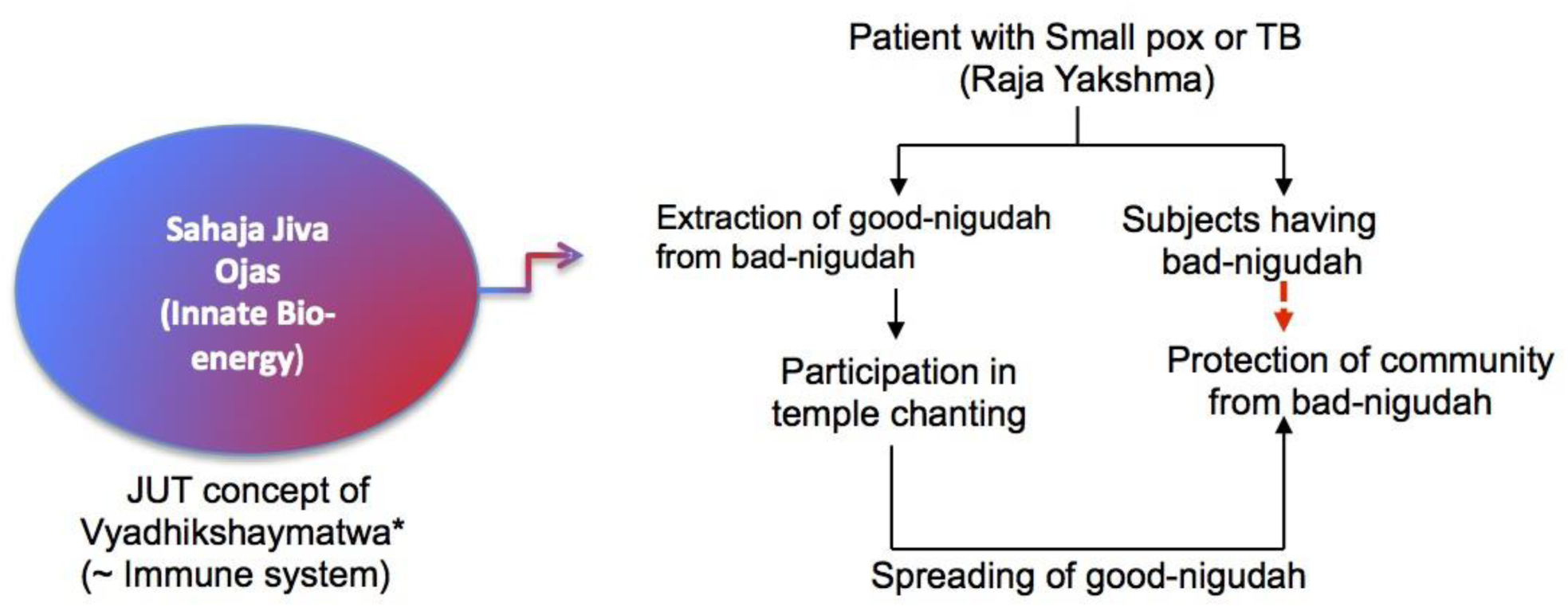
JUT’s primary focus has been to treat patients suffering with disease caused by Nigudahkrimis (invisible krimis or microscopic life form as per Ayurveda, also known as nigoda in Jain sacred texts). Nigudah krimis may cause disease, when the number of good (Upakari or sahaja or suad) nigudah declines, and bad (Apakari or piscacha) nigudah increases in the body because of the weakness in the Nigudah-immunity. JUT-based traditional medicine man (tantric ojah) try to tilt the balance in a sick person by extracting good-nigudah from bad-nigudah by activating a subtle defense mechanism of the body, the Sahaja Jiva Ojaskshyamatwa (Nigudah-immunity), a part of the Sahasa-ojash or Avatar Kosha (details in, page 16, Reference 17). This idea of Nigudah-Ojaskshayamatva or Nigudah-immunity is similar to Vyadhikshaymatwa or Ojaskshyamatwa, a basic concept of Ayurvedic immune defense mechanism (34; 35; 36). The increase of good-nigudah in the body lead to speedy recovery. JUT healers or Ojah employ the recovered person as a source of aerosol-inoculation to spread good-nigudah in the community (the treatment method was originally used against small pox). In this manner, the bad-nigudah of sick patients (broken red arrow) cannot spread into the community. Thus, JUT based tantric healers try to enhance Sahaja Jiva Ojas, or innate bio-energy of patient’s body (details in, page 8-19, Reference 17).

#### Satvata tarka, and Prajna Agan (Knowledge emergence)

JUT uses a tarka method as a part of the knowledge emergence. “Tarka” (suppositional reasoning) is a well-established investigation method for new knowledge generation in ancient Indian philosophical system (37). Tarka emphasizes on the social dimension of knowledge, where resolving controversy is reined by reasoning rather than the sources of knowledge. Paradigmatically, tarka is called for in order to establish a presumption of truth in favor of one thesis against a rival thesis, both of which are supported by a putative source of knowledge. The thesis and the counter thesis both are backed up by apparent knowledge sources such as genuine inferences (the most common situation) or competing perceptual or testimonial evidences. One scholar tries to refute the reasoning of the competing scholar and try to establish the knowledge validity of his/her own thesis by identifying weakness in opponent’s thesis. These weaknesses include anista-prasanga (unacceptable consequences) or breaks in intellectual norms in the rival’s knowledge. It is presumed that such tarka event, which can be compared to modern world peer-review concept, may enhance the overall knowledge of both the groups leading to increase of overall knowledge generation in the society including the knowledge-emergence process of IKS (14).

The tarka based reasoning spread throughout ancient India, and was known as Anviksiki during the composition of Nyaya Sutra by Gotama (500 BC), and then probably spread throughout India (97, 110–112) as evident by the presence of Tarka culture in Kamarupa (present day Assam). The tarka was probably adapted by various local institutions through the process of IKS by knowledge generation (Chapter 5, Reference 17).

**Figure 2:**
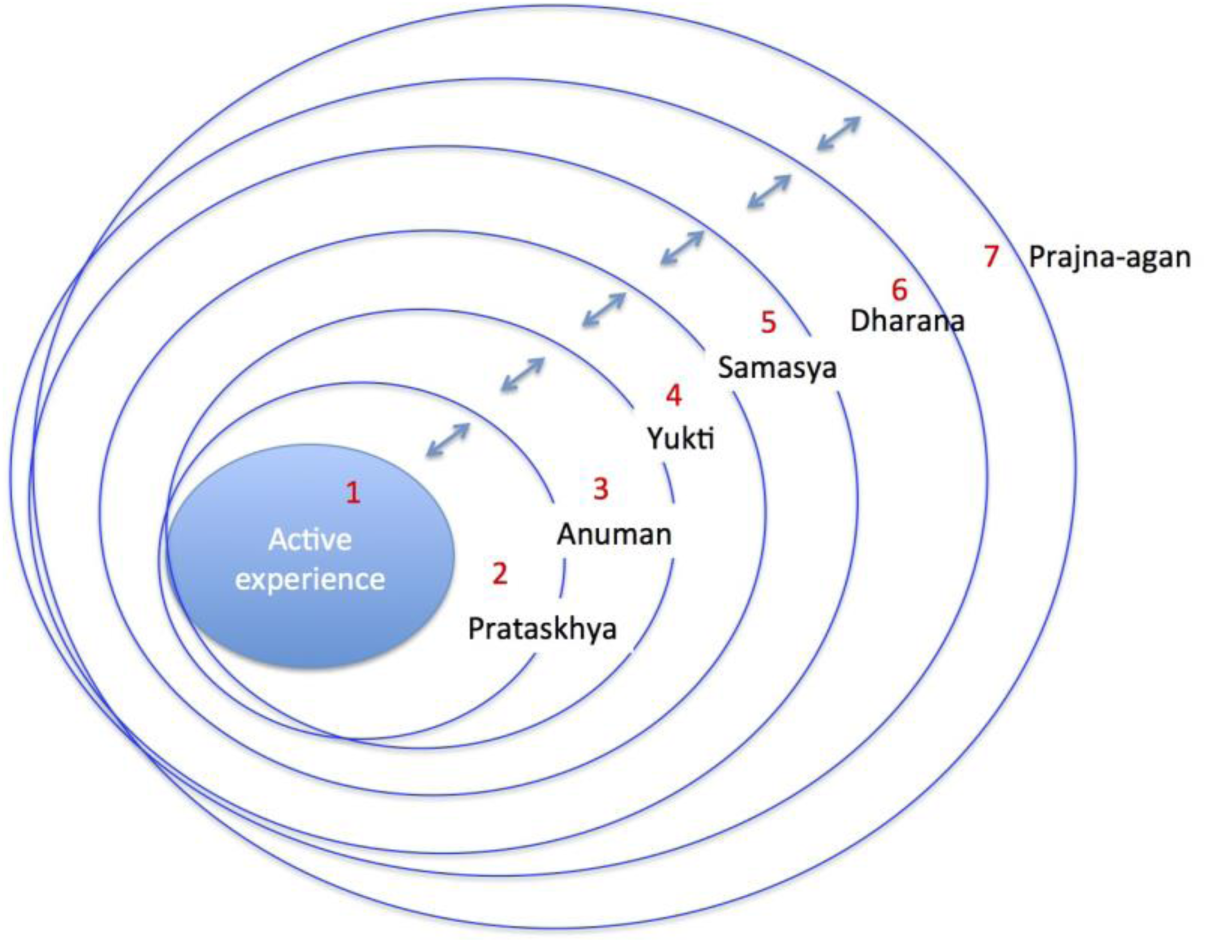
The Guru Asana Vyuha component of Satvata Tarka. The dynamic interaction among the circles is known as Satvata sanjog (realistic connection). Adapted from Chapter 2, Reference 17

Two co-authors of this manuscript, Jagat Ghora (JG) and Krishna Ram Das (KRD) used an unique JUT based tarka system; they referred to an ancient Satvata (examination of truth) anviksiki tarka method. JG adapted this tarka for his tantric-healing practice (Chapter 1.4, Reference 17); Krishna Ram Das (KRD) used a similar anviksiki tarka based method for poetry writing (page 53, Reference 98, and 25). According to KRD, the Satvata tarka was practiced in Hatisatra temple, and had five circles (pancha-cakra) or stages of debate (Figure 2):

1. Observation of a desire (ichha-prataskhya)
2. Inference based on prior knowledge (satvata/purvavat-anuman)
3. Suppositional reasoning (yukti-tarka)
4. Special feature of doubt (samasya-laksana) and
5. Examination of a new idea (parkisa-dharana)

Some of these five elements were part of an ancient system of knowledge generation through reasoning (page 2 and 32, reference 37), whereas the other elements are the product of local transformation of knowledge system as predicted by local adaptation of indigenous knowledge system (14, 99).

These five elements or circles of tarka are initiated by two foundational element of tarka (Figure 2):

1. Activity based postulation (pravrtti-arthapati) and
2. Active experience (Vijana).

These seven concentric circles are represented by one big idea/view/circle (Figure 2), and known as guru-asana vyuha (the concentric seat of a guru).

**Figure 3:**
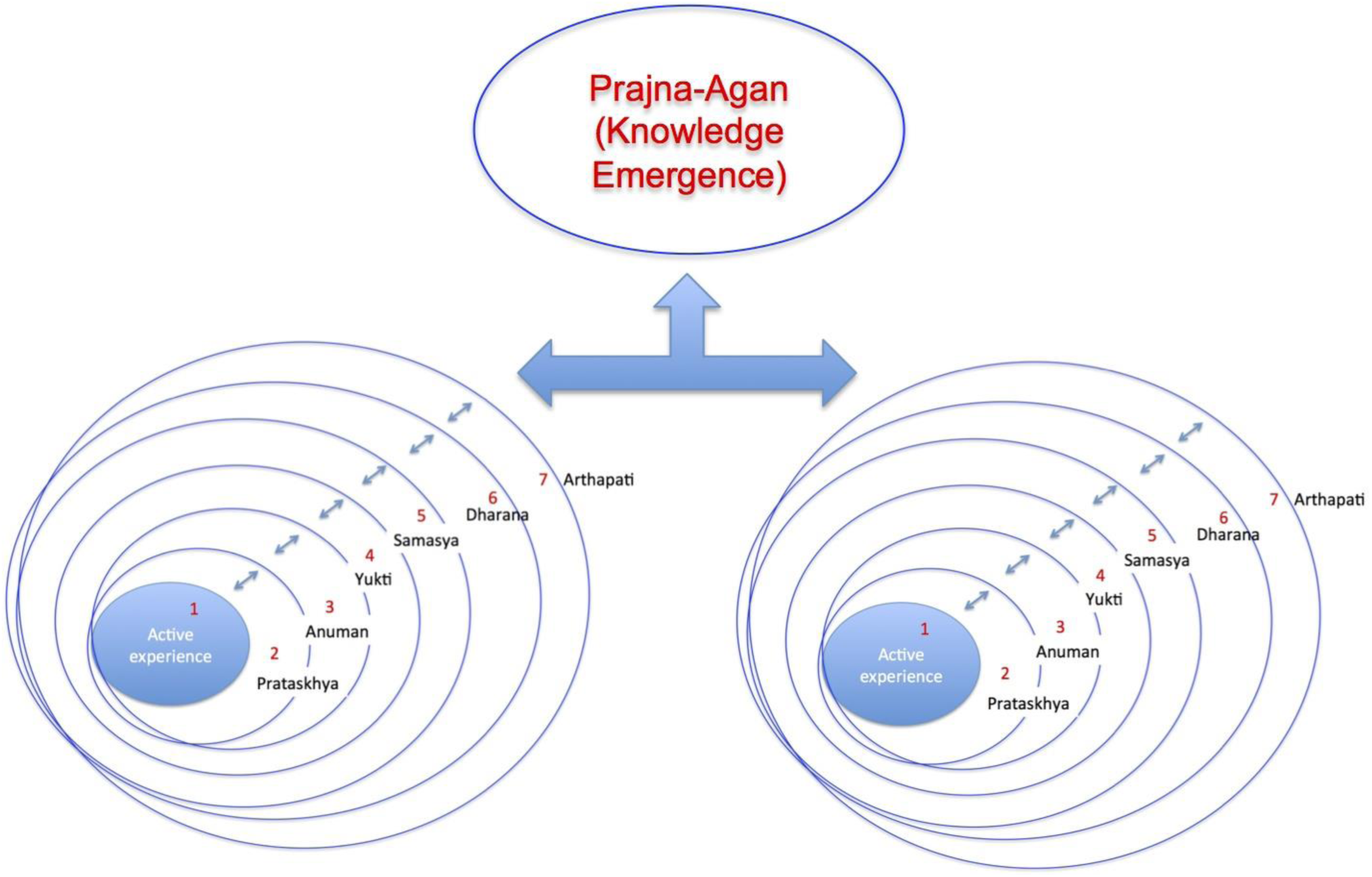
Prajna-Agan: the process of knowledge emergence through the dialectic process of Tarka between two opposite ideas. Adapted from Chapter 2, Reference 17.

In the Satvata tarka process, one such idea/circle engage in argument with an opposite idea/circle through a structured process, where each of the idea go through “parnima” (transformation) to sharpen each other’s “yukti” (reasoning). This process of dynamic interaction, “satvata sanjog” between the two opposite ideas lead to “prajna agan” i.e. knowledge emergence, a component of IKS (14); Figure 3-4. Taken together, Satvata tarka is dialectic method of reasoning, probably having its root in ancient Kamarupa (Chapter 2, Reference 17).

**Figure 4:**
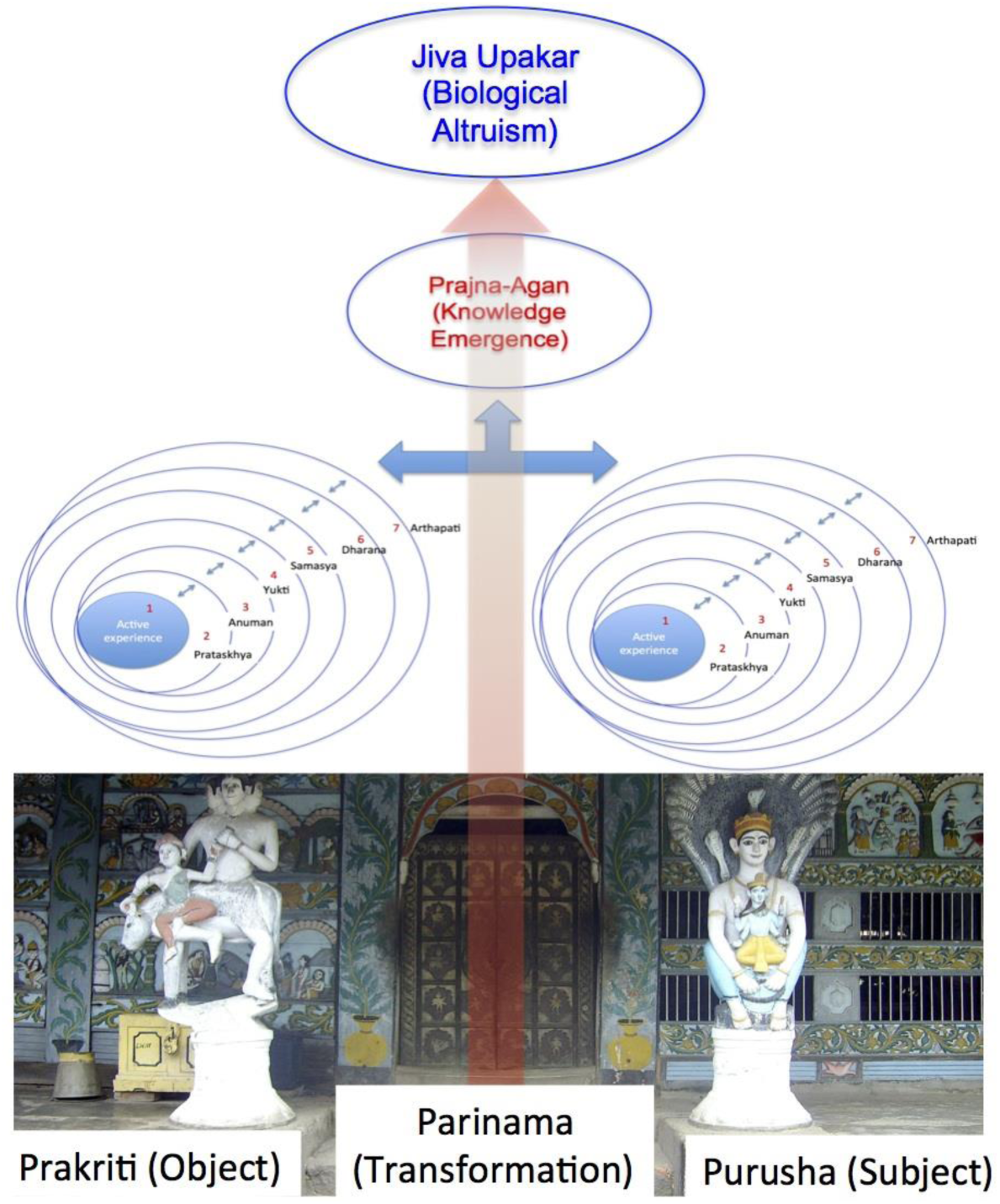
Schematic representation of Jiva Upakar Tantra based metaphysis of knowledge emergence. The right interaction between our physical body (Prakriti) and self (subtle body of Purusha) lead to parinama (transformation) of Ojas (bio-energy, shown as the reddish arrow going upward; the door of Vyadhikshamatva or body defense) to Jiva Upakar (biological altruism) Shakti (strength/power).

Taken together, JUT or Vedic Altruism is an Ayurvedic concept of Vyadhikshamatva that probably emerged as an IKS in the Hatisatra temple following centuries of local Ayurvedic practice led by Ojhas (traditional tantric healers) (Chapter 2 and 3, Reference 17).

### Example of Prajna-agan (Knowledge emergence): First example: FGD session to design a contact investigation study (Figure 1-3 main text)

As depicted in Figure 1 in this note, we were able to formulate the metaphysics behind JG’s good-nigudah based aerosol inoculation against PTB. The epistemology of this metaphysics, the “satvatatarka” or dialectic method of reasoning is based on Figure 2-4 in this note. During this tarka session, BD expressed his concern about JG’s practice of aerosol inoculation, as it may encourage *Mtb* transmission in the community. At the same time, BD wanted to know more about the metaphysics that underlies JG’s healing practice. Hence, JG, KRD, and five other participants participated in a FGD session in the Hatisatra temple complex (Figure 4).

**Figure 4:**
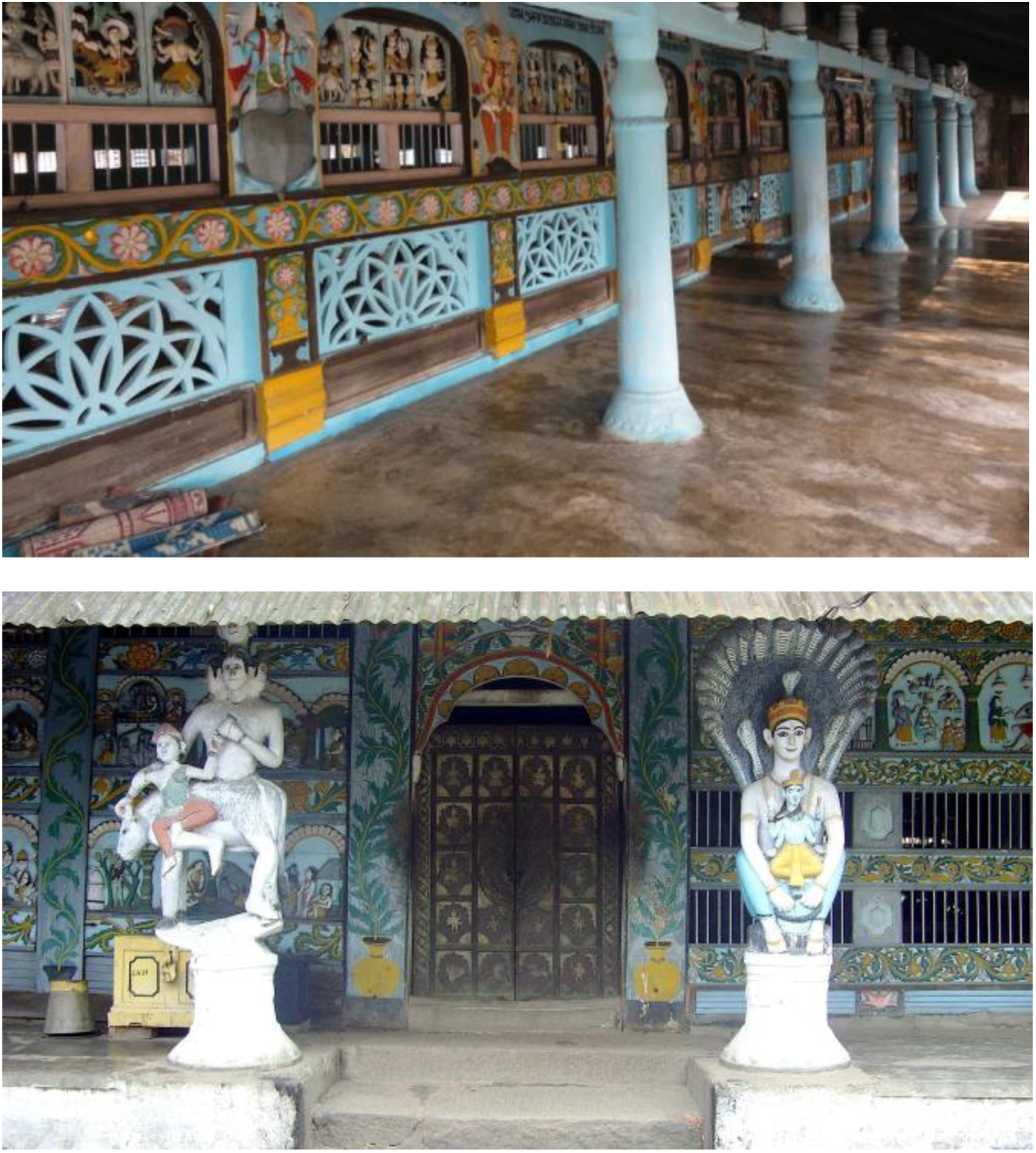
Hatisatra temple (top photo, the courtyard of the temple; bottom photo, the northern gate of the temple), where the first and second FGD session of Satvata tarka took place between BD, and JG/KRD in 1994, and BD/JG in 1998. During the tarka session, JG repeatedly showed the north door of the temple (bottom photo) as a representative image of Satvata tarka. The two deities (Shiva or Prakriti on the left, and Vishnu or Purusha on the right) represent two circles of the tarka (Figure 3 & 4). The door in the middle with the depiction of ten Avatars of Vishnu represents the Prajna-Agan component of the tarka. Since the interaction results in the emergence of Avatars, who perform altruistic work in society, the door represents a physical embodiment of Jiva Upakar vada (theory of biological altruism). Thus, according to JG/KRD, Satvata Tarka is the knowledge-generation component of JivaUpkar Vad, a possible derivation of Vedanta’s Parinamvada (the cause and effect dualism of Vedanta). Interestingly, Parniamvada concept has similarity with Western dialectic (113).

### Second example: Visualization of good nigoda through Satvata Tarka

After completion of the 15 years of initial contact investigation study (1994–2009), we did the second Satvata tarka session to discuss the study results. Dayal Krishna Bora (DKB), an eminent vedic scholar agreed to represent JG and KRD, as both of them passed away. DKB was familiar with JG and KRD’s Satvata tarka method, as the trios were involved in researching on Jiva Upakar Tantra (page 53, 428-530, Reference 98). Thus, DKB represented the good Nigoda-hypothesis side, whereas BD represented the modern Immunology side (two circles of tarka, Figure 2A). The tarka started with initial enumeration (udeshya): to find out why chanting showed apparent protection from PTB. Then, the tarka process started with the postulation (pravritti-arthapati) stage. BD argued from the Immunology point of view (the Immunology circle) that apparent protection was due to latent *Mtb* infection acquired during chanting; there body developed immunity against the pathogen. This idea was opposed by DKB, who provided argument based on Jiva Upakar Tantra (JUT): the apparent protection was due to spread of good nigoda. Then, DKB moved to the Yukti-tarka circle, and identified the element of anista-prasanga **yukti** (reasoning based on unwanted consequence) in BD’s presumption. DKB stated that BD’s conclusion that devotee may be infected by the pathogen is an unwanted effect of BD’s initial observation that SN-PTB subjects do not show bacteria in sputum; without bacteria in their sputum, how could they spread the pathogen to others? BD replied that 30% of SN-PTB subjects convert to smear-positive subjects as the disease progress. So, it is possible that 3 of the smear-negative chanting devotees of the study eventually became smear positive and infected the rest, who received low-dose of bacteria enough to trigger immunity to resist bacterial growth in lung. Those who did not attend Kirtan did not get infected with a low dose bacterium, and therefore, when they were exposed to high dose bacteria by smear-positive people in the community, they developed PTB. However, DKB reached an alternative explanation by applying the **purvavat-anuman** (unperceived effect from a perceived cause) (37, 97). DKB inferred that good nigoda spreading from the SN-PTB subjects during Kirtan chanting protected the rest of the devotees from TB by inducing immunity. To this explanation, BD applied **samasya-laksana** (doubtful feature) and instated that good nigoda is an imaginary thing, and therefore, cannot be a factor for reasoning. To this argument, DKB applied **satavata-anuman** (inference based on prior knowledge) to conceptualize the physical feature of a good-nigoda. DKB referred to an ancient law of duality (dvaita-vedanta), where it is possible for good to emerge from bad through the process of protibimbavada (law of reflection). The theory was proposed by an 8^th^ century vedantic scholar Padmapada that certain effects may be the mirror image of a cause (110). DKB asked BD to draw the general features (laksana) of the TB causing bacteria in a piece of paper. BD drew a seed (DNA and cytoplasm) inside a sack (plasma membrane), Figure 4B, main text. DKB then applied the law of reflection to draw a similar sack with part of the seed inside, and termed it as good-nigoda. DKB then invoked the observation of a desire (**ichha-prataskhya**) and cited the example of Dadhicchi (an ancient sage who donated his own bone for greater cause of social benefit) to infer that some good chemical exist in the bacteria, which will be present in the seed of good-nigoda. DKB then asked BD to come up with any known features by medical science that can resemble the image of good-nigoda (Figure 4B, main text). To this question, BD replied that bacteria-infected cells of our body might be able to capture some antigens from the bacteria, and put them inside a vesicle and secrete them outside; such vesicles are known as extracellular vesicles (EVs). DKB then invoked the observation of a desire (**ichha-prataskhya**) and postulation (arthapati) elements of the tarka to insist that good-nigoda can be observed in the aerosol of smear-negative subjects. Both DKR and BD then invoked the last element of the tarka, **pariksha-dharana** or the examination of a new idea, and concluded that prajna-agan or knowledge emergence happened, as we obtained an experimental idea to directly test the existence of good-nigoda in aerosols. Thus, we drew the image of the good-nigoda (Figure 4B, main text), and hypothesize that an extracellular vesicle-containing antigen derived from *Mtb* may serve as the model for good nigoda. In this manner, Satvata tarka was performed during FGD sessions with DKB.

